# Quantifying Excitation–Inhibition Harmony and Its Breakdown during Epileptogenesis in a Spiking Neural Network of the Hippocampal CA1

**DOI:** 10.64898/2026.11.02.763777

**Authors:** Sang-Yoon Kim, Woochang Lim

## Abstract

Epileptogenesis is often linked to disruptions in excitation–inhibition (E-I) balance, yet its quantitative characterization remains limited. Understanding and quantifying this E-I balance could enable earlier diagnosis and targeted treatment of epilepsy. We investigate competitive E–I harmony and its breakdown using a biologically grounded spiking neural network model of hippocampal CA1 that incorporates two structural pathological mechanisms: pyramidal cell (PC) axonal sprouting and loss of inhibitory interneurons. As recurrent excitatory connectivity increases via PC axonal sprouting or inhibitory interneuron populations decline through interneuron loss, the network undergoes sequential transitions from a healthy phase (HP) to an interictal pathological phase (IIP) and finally to an ictal pathological phase (IP). At the input level, we introduce the competition degree, *C*_*d*_ (= *S*_*E*_*/S*_*I*_), as the ratio of the strengths of excitatory (*S*_*E*_) to inhibitory (*S*_*I*_) synaptic inputs to PCs, to quantify their competitive E-I harmony. As an input-side indicator, *C*_*d*_ remains near 2.6 in HP, exhibits a discontinuous, abrupt jump to ~ 60 at the HP → IIP transition and rises further thereafter. At the output level, the corresponding output-side measure, the population-averaged mean firing rate (MFR) of PCs, 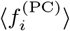, also exhibits a discontinuous, abrupt jump from ~ 4 to ~ 200 Hz at the HP → IIP transition and rises further thereafter, indicating progressive pathological hyperexcitability. Together, these results establish a cause-and-effect framework linking the input-side indicator *C*_*d*_ (cause) to the output-side measure 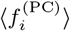 (effect). Accordingly, *C*_*d*_ may serve as a potential quantitative biomarker of competitive E–I harmony and its breakdown, providing a computational causal framework that links structural pathological mechanisms to epileptogenic state transitions and offering potential for translation into diagnostic tools and therapeutic applications.

## I. INTRODUCTION

Epilepsy is a chronic neurological disorder marked by recurrent, unprovoked seizures, arising from abnormal hyperexcitable neuronal activity [1–5]. A widely accepted mechanism underlying seizure generation is the disruption of the so-called “E-I balance” in neural circuits [6–17]. While this framework has guided decades of research, it typically remains qualitative, focusing on whether excitation exceeds inhibition rather than providing a quantitative measure of their interaction.

The classical notion of E–I balance suggests equality of excitation and inhibition, but healthy networks typically exhibit excitatory dominance dynamically controlled by inhibition. We therefore introduce the concept of competitive E–I harmony, in which excitation provides the primary drive for physiological function, while inhibition controls excitation. Their competitive yet cooperative interplay sustains stable network function.

Previously, we introduced the “competition degree” (*C*_*d*_) in the basal ganglia to quantify the interplay between the direct “Go” and indirect “No-Go” pathways, which exert opposing influences on the inhibitory output nucleus, the substantia nigra pars reticulata (SNr) [18, 19]. The competition degree *C*_*d*_ is given by the ratio of the strengths of the Go pathway to the No-Go pathway. In healthy motor control, *C*_*d*_ ~2.8, reflecting a slight dominance of the Go pathway and stable motor behavior, whereas deviations of_*d*_ from this healthy value reflect competitive disharmony, as in Parkinson’s or Huntington’s disease. Here, we extend this concept to the hippocampal CA1 by defining *C*_*d*_ = *S*_*E*_*/ S*_*I*_, the ratio of the strengths of excitatory (*S*_*E*_) to inhibitory (*S*_*I*_) synaptic inputs to pyramidal cells (PCs). Healthy physiological states exhibit *C*_*d*_ ~2.6, reflecting E–I harmony and stable function, whereas deviations of *C*_*d*_ from this healthy value reflect the breakdown of E–I harmony. This competition-degree approach offers new insights into the dynamical basis of epileptogenesis, the process by which a normal brain becomes susceptible to recurrent seizures [20].

Disruption of E–I harmony through excessive excitation or loss of inhibition has long been recognized as a fundamental mechanism in epileptogenesis [6–17]. Among the hippocampal subregions, CA1 is particularly susceptible to seizure generation and propagation [21– 25], making it a key target in the study of temporal lobe epilepsy [26–32]. Two key structural alterations in CA1 have been implicated in epileptogenesis: (i) axonal sprouting of PCs, enhancing recurrent excitatory connectivity [33–40], and (ii) loss of inhibitory interneurons, such as basket cells (BCs) and oriens-lacunosum moleculare (O–LM) cells [40–45]. While these phenomena are well documented, most studies have focused primarily on the output-side effects, such as pathological firing activities, rather than quantitatively characterizing the input-side E–I competition that gives rise to these effects. Thus, how these structural alterations reshape excitatory and inhibitory synaptic inputs to PCs and thereby drive output-side pathological firing activity and epileptogenic state transitions remains to be quantitatively characterized.

Computational modeling of CA1 has been applied at various scales, from mean-field approaches to spiking neural networks (SNNs), reproducing features such as interictal discharges and ictal-like events [46–65]. However, most studies examined isolated pathological states or specific mechanisms, without demonstrating the full progression from healthy to interictal to ictal dynamics or quantitatively linking input-side changes to their output-side effects. Consequently, the successive transitions of epileptogenesis and their underlying cause-and-effect relationships remain underexplored within a single CA1 framework.

To address this gap, we develop a biologically inspired SNN of hippocampal CA1 incorporating excitatory PCs and inhibitory BC and O–LM cells. We investigate two pathological mechanisms: (i) axonal sprouting of PCs, modeled by increasing the recurrent excitatory connection probability *P*_*c*_ among PCs, and (ii) loss of inhibitory interneurons, modeled by reducing BC and O–LM populations. Numerical study of the CA1 SNN reveals three successive phases as either mechanism intensifies: a healthy phase (HP) with irregular low-rate output-side PC firing, an interictal pathological phase (IIP) with synchronous output-side PC discharges, and an ictal pathological phase (IP) with irregular high-rate output-side PC firing.

To quantify these transitions characterized by the output-side firing activities of PCs, we introduce the competition degree *C*_*d*_ (= *S*_*E*_*/ S*_*I*_) as an input-side indicator of E-I competition. In HP, *C*_*d*_ ~2.6 reflects E-I harmony, whereas at the HP → IIP transition,_*d*_ exhibits a sharp, discontinuous jump to *C*_*d*_ ~60, followed by a gradual rise through IIP and IP. Correspondingly, the output-side measure, the population-averaged mean firing rate (MFR) of PCs, 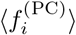, exhibits an abrupt, discontinuous jump from ~4 to ~200 Hz at the HP IIP transition and rises further thereafter, indicating progressive pathological hyperexcitability. Thus, the discontinuous jump in_*d*_ provides a quantitative input-side signature of the HP →IIP transition and is directly associated with the corresponding abrupt change in the output-side measure 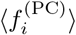

Although the present study is computational, *C*_*d*_ has potential translational relevance. Because *C*_*d*_ is defined in terms of excitatory and inhibitory synaptic inputs, it may be estimated using invasive electrophysiological recordings [66, 67] or advanced imaging approaches to synaptic function [68, 69], offering a potential biomarker for early detection, disease monitoring, and assessment of therapeutic interventions. Moreover, because disruptions in E–I harmony are implicated in other neurological and psychiatric conditions [70–77], the causal framework we present may extend beyond epilepsy to inform broader clinical strategies.

In summary, this work provides a quantitative causal framework for the breakdown of E-I harmony during epileptogenesis, linking the input-side indicator *C*_*d*_ to the corresponding output-side measure, the population-averaged MFR of PCs, 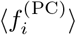, and offering potential avenues for early detection and targeted interventions.

Finally, the remainder of this paper is organized as follows. Section II describes the construction of the hippocampal CA1 SNN. Section III presents a PC-focused, causal-framework-based analysis of E–I harmony and its breakdown during epileptogenesis. Section IV provides a summary and discussion. The Appendices provide details of the CA1 SNN and the firing activities of interneurons during epileptogenesis.

## II. BIOLOGICALLY INSPIRED SPIKING NEURAL NETWORK OF THE HIPPOCAMPAL CA1

In this section, we briefly describe our biologically inspired SNN of the hippocampal CA1, with details provided in Appendices A and B. Based on the anatomical and physiological properties of CA1 described in [56– 59, 78–81], we construct our CA1 SNN by determining the cell population sizes and synaptic connection probabilities. As single-cell models for PCs, BCs, and O-LM cells, we employ the Izhikevich neuron models [82– 85] and adjust their intrinsic parameters to match the experimentally obtained *f* − *I* curves [86–88]. We also incorporate receptor-mediated synaptic current models, whose detailed formulations are given in Appendix B.

### A. Framework of the CA1 SNN

Figure 1 shows the box diagram for our CA1 SNN. It consists of excitatory PCs and two types of inhibitory interneurons, BCs and O-LM cells. There are two external sources, entorhinal cortex (EC) and CA3. The EC provides excitatory input to PCs via the perforant path (PP), while the CA3 sends excitatory input to both PCs and BCs via Schaffer collaterals (SC). The principal PCs send excitatory outputs to the subiculum (S) and EC.

**FIG. 1:**
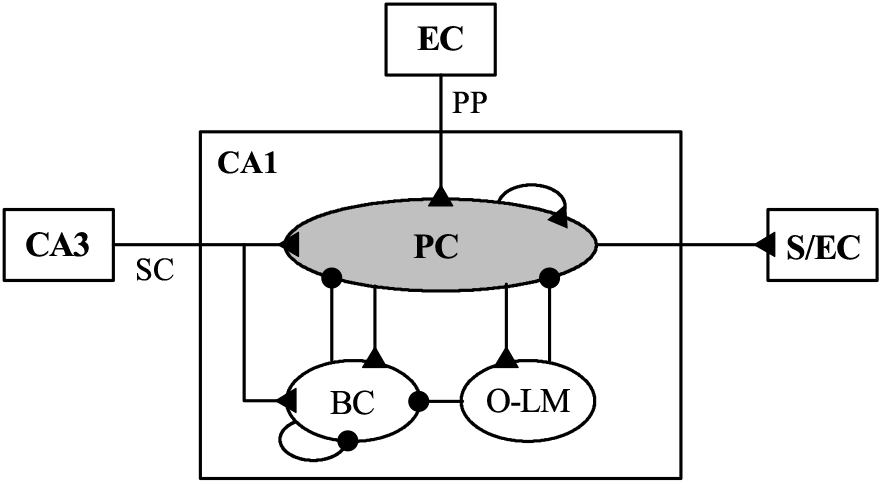
Box diagram of the spiking neural network of the hippocampal CA1. Excitatory and inhibitory connections are denoted by lines with triangles and circles, respectively. Pyramidal cells (PCs) receive excitatory inputs from the entorhinal cortex (EC) via the perforant path (PP) and from CA3 via Schaffer collaterals (SC). In the CA1, we consider two types of inhibitory interneurons: basket cells (BCs) and oriens-lacunosum moleculare (O-LM) cells. PCs send excitatory outputs to the subiculum (S) and EC.

BCs provide feedforward (CA3 → BC → PC) and local feedback (BC ↔ PC) inhibition to PCs. BCs send inhibitory projections to the somata of PCs, thereby controlling their firing activity. In contrast, O-LM cells provide only feedback (O-LM ↔PC) inhibition to the dendrites of PCs, controlling excitatory inputs to PCs from EC and CA3. O-LM cells also control the firing activity of BCs (O-LM → BC), but there is no reciprocal projection from BCs to O-LM cells (BC → O-LM).

We note that PCs receive excitatory (E) synaptic inputs from three sources (EC, CA3, and recurrent PC projections) and inhibitory (I) synaptic inputs from two sources (BCs and O-LM cells). In the tonic resting state, E-I balance is achieved, whereas in the phasic healthy active state, E-I harmony is maintained. When E-I harmony breaks down, pathological states emerge during epileptogenesis. These states and the breakdown of E-I harmony are quantitatively analyzed in Sec. III.

Based on the anatomical data given in [78–81], we choose the numbers of constituent cells (PCs, BCs, and O-LM cells) in the CA1, PCs in layer III of the EC, and PCs in the CA3. The number of PCs in CA1 (*N*_PC_) corresponds to about 89 % of the total number of CA1 cells (*N*_tot_). There are various types of interneurons in the CA1, whose total number (*N*_IN_) corresponds to about 11 % of *N*_tot_. Thus, *N*_PC_: *N*_IN_ ≃8 : 1. Here, we consider two types of interneurons, BCs and O-LM cells, among the diverse types of interneurons. We set the number of O-LM cells to *N*_O*−*LM_ = 100 and determine the numbers of the other cells using the population-size ratios *N*_EC_: *N*_CA3_: *N*_PC_: *N*_BC_: *N*_O*−*LM_ ≃28.4 : 23.47 : 35.3 : 3.4 : 1, where *N*_EC_ and *N*_CA3_ denote the numbers of PCs in the EC and CA3, respectively. Accordingly, we obtain *N*_EC_ = 2840, *N*_CA3_ = 2347, *N*_PC_ = 3530, *N*_BC_ = 340, and *N*_O*−*LM_ = 100, as summarized in Table I.

**TABLE I:** Numbers of cells in the hippocampal CA1 SNN and its external input populations.

|  |  |  |
|---|---|---|
| *N*<sub>EC</sub> (PCs in EC layer III) |  | 2,840 |
| *N*<sub>CA3</sub> (PCs in CA3) |  | 2,347 |
| CA1 | *N*<sub>PC</sub> (PCs) | 3,530 |
|  | *N*<sub>BC</sub> (Feedforward Basket cells) | 340 |
|  | *N*<sub>O−LM</sub> (Feedback O-LM cells) | 100 |

Next, we choose the connection probabilities between cells in the CA1 SNN, based on [57–59]. From outside the CA1, the EC sends random excitatory projections to PCs via PP with the connection probability 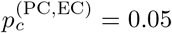. Also, the CA3 provides excitatory inputs randomly to PCs and BCs via SC with the connection probabilities 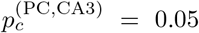 and 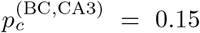, respectively. Within the CA1, random synaptic connections between constituent cells are considered; random recurrent connections between PCs and between BCs are also considered. Table II shows the synaptic connection probabilities 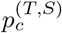 from a presynaptic cell in the source population (*S*) to a postsynaptic cell in the target population (*T*).

**TABLE II:**
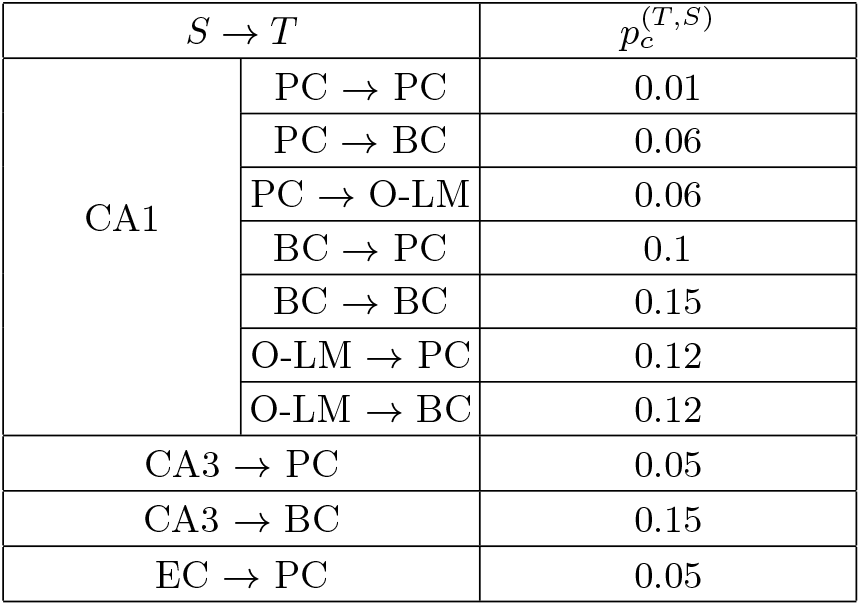
Synaptic connection probabilities 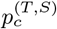 from a presynaptic cell in the source population (*S*) to a postsynaptic cell in the target population (*T*).

| *S* → *T* |  | *p*<sub>*c*</sub><sup>(*T*,*S*)</sup> |
|---|---|---|
| CA1 | PC → PC | 0.01 |
|  | PC → BC | 0.06 |
|  | PC → O-LM | 0.06 |
|  | BC → PC | 0.1 |
|  | BC → BC | 0.15 |
|  | O-LM → PC | 0.12 |
|  | O-LM → BC | 0.12 |
| CA3 → PC |  | 0.05 |
| CA3 → BC |  | 0.15 |
| EC → PC |  | 0.05 |

Finally, external inputs from EC and CA3 are modeled using *N*_EC_ and *N*_CA3_ independent Poisson spike trains with spike rates *f*_EC_ and *f*_CA3_, respectively. In the tonic resting state, *f*_EC_ = 3 Hz and *f*_CA3_ = 1 Hz, while in the phasic healthy active state, *f*_EC_ = 8 Hz and *f*_CA3_ = 3 Hz [57, 89–94].

### B. Single Neuron Models and Synaptic Currents

For the CA1 SNN, we employ the biologically plausible and computationally efficient Izhikevich spiking neuron model [82–85], as in our previous studies [18, 19]. The Izhikevich model reproduces neuronal firing dynamics through appropriate tuning of its intrinsic parameters. The CA1 SNN consists of three neuronal populations: PCs, BCs, and O-LM cells. The state of a cell in each population is characterized by two dynamical variables of the Izhikevich model: the membrane potential *v* and the slow recovery variable *u*. Three types of input currents, *I*_*bg*_ (background current), *I*_*syn*_ (synaptic current), and *I*_*stim*_ (injected stimulation current), govern the time evolution of *v* and *u*.

Here, *I*_*bg*_ represents the background current originating from external regions not explicitly included in the model, while *I*_*stim*_ denotes the injected stimulation DC current. When the membrane potential reaches the spike cutoff threshold, the neuron fires a spike, after which the membrane potential *v* and the recovery variable *u* are reset.

Detailed descriptions of the Izhikevich neuron models for PCs, BCs, and O-LM cells are presented in Appendix A. The Izhikevich neuron model for each cell has nine intrinsic parameters, whose values are listed in Table III in Appendix A. These values are based on the physiological properties of CA1 cells [86–88].

**TABLE III:**
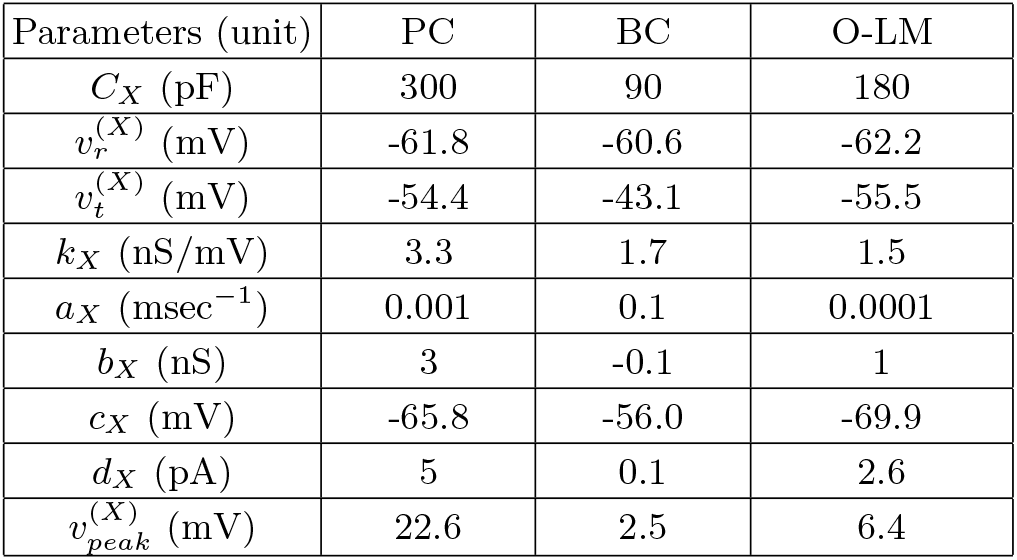
Values of the nine intrinsic parameters of the single-neuron models for PCs, BCs, and O-LM cells.

Next, we consider the synaptic currents *I*_*syn*_ into the CA1 cells. We follow the standard formalism for *I*_*syn*_ [18, 19, 95–100]. Details are provided in Appendix B. Here, we consider two types of excitatory synaptic currents mediated by AMPA and NMDA receptors. AMPA receptors mediate fast excitatory responses, while NMDA receptors mediate slower excitatory responses subject to voltage-dependent Mg^2+^ block.

We also consider two types of inhibitory synaptic currents mediated by GABA_A_ and GABA_B_ receptors. GABA_A_ receptors mediate rapid inhibition, whereas GABA_B_ receptors produce slower, prolonged inhibition. The synaptic conductance associated with each synaptic current can be approximated by the product of the maximum conductance per synapse, the average number of afferent synapses, and the fraction of open postsynaptic ion channels. Postsynaptic ion channels open through the binding of neurotransmitters to receptors in the target population. The temporal evolution of the fraction of open ion channels is described by a sum of double-exponential functions over presynaptic spikes, controlled by the synaptic rise time constant, the synaptic decay time constant, and the synaptic latency time constant. The values of the synaptic parameters, including the maximum synaptic conductance, the synaptic rise time constant, the synaptic decay time constant, the synaptic latency time constant, and the synaptic reversal potential, are given in Table V in Appendix B [56–59].

## III. CAUSAL-FRAMEWORK-BASED ANALYSIS OF E-I HARMONY AND ITS BREAKDOWN DURING EPILEPTOGENESIS

Competitive E-I harmony is essential for maintaining stable physiological activity in the hippocampal CA1. E-I harmony emerges from competitive yet cooperative interactions between excitatory and inhibitory synaptic inputs to PCs. Structural pathological alterations, such as PC axonal sprouting and loss of inhibitory interneurons, disrupt E-I harmony, causing a discontinuous abrupt transition from HP (healthy phase) to IIP (interictal phase), followed by further progression to IP (ictal phase), leading to pathological hyperexcitability and seizures.

To quantitatively characterize competitive E-I harmony and its breakdown, we employ the competition degree *C*_*d*_ (=*S*_*E*_*/S*_*I*_), defined as the ratio of the strengths of excitatory (*S*_*E*_) to inhibitory (*S*_*I*_) synaptic inputs to PCs. This serves as an input-side indicator (cause). As the corresponding output-side measure (effect), we consider the population-averaged MFR of PCs, 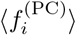. At the HP → IIP transition, *C*_*d*_ exhibits a discontinuous, abrupt jump from ~ 2.6 to ~ 60, accompanied by a corresponding jump in 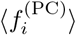 from ~ 4 to ~ 200 Hz. Thereafter, both quantities increase progressively through IIP and IP, revealing a direct correspondence between the breakdown of input-side E-I harmony and the emergence and progression of output-side pathological hyperexcitability.

Within this causal framework, we focus on PCs and systematically investigate (1) E-I harmony in HP and (2) the breakdown of E-I harmony and emergence of pathological states caused independently by PC axonal sprouting and interneuron loss. We examine how the breakdown of input-side E-I harmony, quantified by *C*_*d*_, causes changes in the corresponding output-side firing activities of PCs during epileptogenesis. In addition to the PC-focused analysis in Sec. III, we also examine the firing activities of interneurons during epileptogenesis in Appendix C.

### A. Healthy States: Tonic Resting State (E-I Balance) vs. Phasic Active State (E-I Harmony)

As explained in Sec. II A, external inputs from EC and CA3 are modeled in terms of *N*_EC_ and *N*_CA3_ independent Poisson spike trains with spike rates *f*_EC_ and *f*_CA3_, respectively. We first consider the case of tonic inputs with *f*_EC_ = 3 Hz and *f*_CA3_ = 1 Hz in the resting state [57, 89–94].

The output-side firing activities of CA1 cells (i.e., PCs, BCs, and O-LM cells) are characterized in terms of population and individual firing activities. Population firing activity of CA1 cells may be well visualized in the raster plot of spikes which is a collection of spike trains of individual CA1 cells. As a collective quantity showing population activities, we employ an IPSR (instantaneous population spike rate) which could be obtained from the raster plot of spikes [101–105]. Each spike in the raster plot is convoluted with a kernel function *K*_*h*_(*t*) to get a smooth estimate of IPSR *R*_*X*_ (*t*) in the *X* population (*X* = PC, BC, and O-LM cell) [106]:

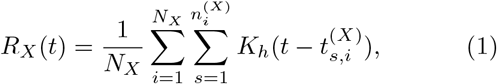

where *N*_*X*_ is the number of the cell, and 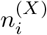 and 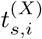 are the total number of spikes and the *s*th spiking time of the *i*th cell, respectively. As the kernel function, we use a Gaussian function of band width *h*:

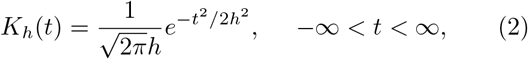

where the band width *h* of *K*_*h*_(*t*) is 20 msec.

Figure 2 shows the output-side population and individual firing activities of the CA1 cells for the tonic resting state. The raster plot of spikes and the IPSR *R*_PC_(*t*) for the PCs are shown in Figs. 2(a1) and 2(a2), respectively. Spikes of PCs are randomly scattered without population synchronization, yielding a noisy, irregular *R*_PC_(*t*) with a low time-averaged value, 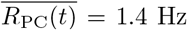. Figure 2(b) shows the histogram of individual MFRs (mean firing rates) 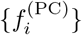 of PCs. In this case, the population-averaged MFR 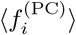 (= 1.4 Hz) is the same as 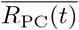 and low; ⟨· · · ⟩ represents population average over all the PCs.

**FIG. 2:**
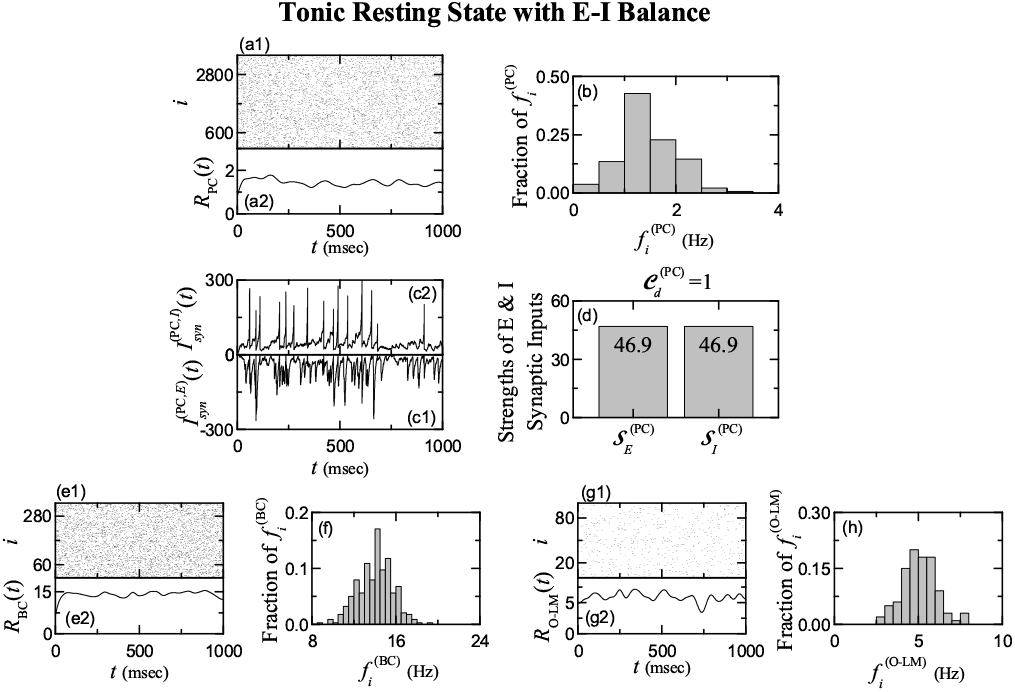
Output-side firing activities (effect) and input-side synaptic quantities (cause) for the tonic resting state with E-I balance (*f*_EC_ = 3 Hz and *f*_CA3_ = 1 Hz). Output-side population and individual firing activities of PCs: (a1) Raster plot of spikes and (a2) IPSR (instantaneous population spike rate) *R*_PC_(*t*) of PCs. (b) Histogram of individual MFRs (mean firing rates) 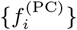 of PCs. Input-side synaptic quantities of PCs: Time series of population-averaged (c1) excitatory (E) 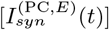 and (c2) inhibitory (I) 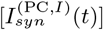 synaptic currents over all PCs. (d) Strengths 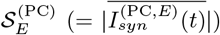 and 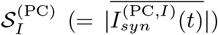 of the population-averaged excitatory and inhibitory synaptic currents. Output-side population and individual firing activities of interneurons: (e1) Raster plot of spikes and (e2) IPSR *R*_BC_(*t*) of BCs. (f) Histogram of individual MFRs 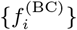 of BCs. (g1) Raster plot of spikes and (g2) IPSR *R*_O*−*LM_(*t*) of O-LM cells. (h) Histogram of individual MFRs 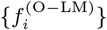 of O-LM cells.

As input-side quantities (cause) underlying the output-side firing activities (effect) of PCs, Figs. 2(c1) and 2(c2) show time series of population-averaged excitatory and inhibitory synaptic currents, 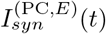 and 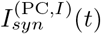, into PCs;

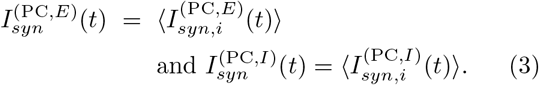

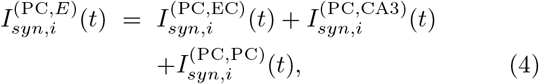

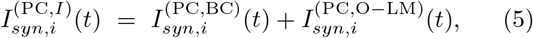

where 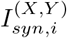 (*t*) represents the synaptic current from the presynaptic source *Y* population to the postsynaptic *i*th cell in the target *X* population.

Strengths of the excitatory and inhibitory synaptic inputs to PCs, *S*_*E*_ and *S*_*I*_, are given by time-averaged magnitudes of the population-averaged excitatory and inhibitory synaptic currents, 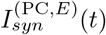 and 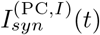:

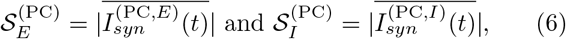

where the overline denotes time-averaging and |···| represents the absolute magnitude. 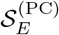 and 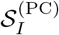 are shown in Fig. 2(d). We note that in the tonic resting state, the excitatory and inhibitory synaptic input strengths are equal, with 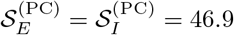.

The output-side firing activity (effect) of the PCs is determined via competition between excitatory and inhibitory synaptic inputs to PCs. To quantify this E-I competition, we introduce the competition degree 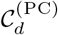 as an input-side indicator (cause), defined as the ratio of 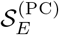 to 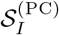:

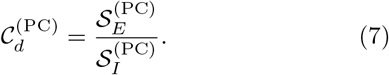

For the tonic resting state, 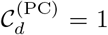, corresponding to E-I balance between the excitatory and inhibitory synaptic inputs to PCs.

We also consider the output-side population and individual firing activities of interneurons (i.e., BCs and O-LM cells) in the tonic resting state. Similar to the case of PCs above, population firing activity is characterized by raster plots of spikes and IPSRs, whereas individual firing activity is quantified using MFRs, as shown in Figs. 2(e1)-2(f) for the BCs and Figs. 2(g1)-2(h) for the O-LM cells.

In both cases of BCs and O-LM cells, spikes are randomly scattered in their raster plots of spikes without population synchronization, as in the case of PCs. Therefore, their IPSRs, *R*_BC_(*t*) and *R*_O*−*LM_(*t*), exhibit noisy, irregular fluctuations around their time average values, 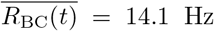 and 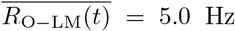, respectively. But, in contrast to the population-averaged MFR 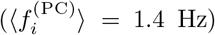 of the PCs, BCs and O-LM cells exhibit firings with higher population-averaged MFRs, 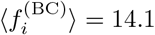 and 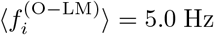, respectively; these population-averaged MFRs are the same as time-averaged IPSRs.

Throughout this paper, the averaging time for the IP-SRs *R*_*X*_ (*t*) (*X* = PC, BC, O−LM), strengths of the excitatory and inhibitory synaptic inputs to PCs, 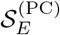 and 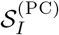, and individual MFRs 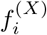 is 10,000 msec. The bin width for the histograms of individual MFRs 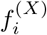 is 0.5 Hz.

Next, we consider the case of phasic inputs of EC and CA3 with *f*_EC_ = 8 Hz and *f*_CA3_ = 3 Hz [57, 89–94], and make quantitative analysis of phasic healthy active state using the input-side competition degree 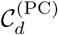. Figure 3 shows the output-side population and individual firing activities of the CA1 cells for the phasic healthy active state.

**FIG. 3:**
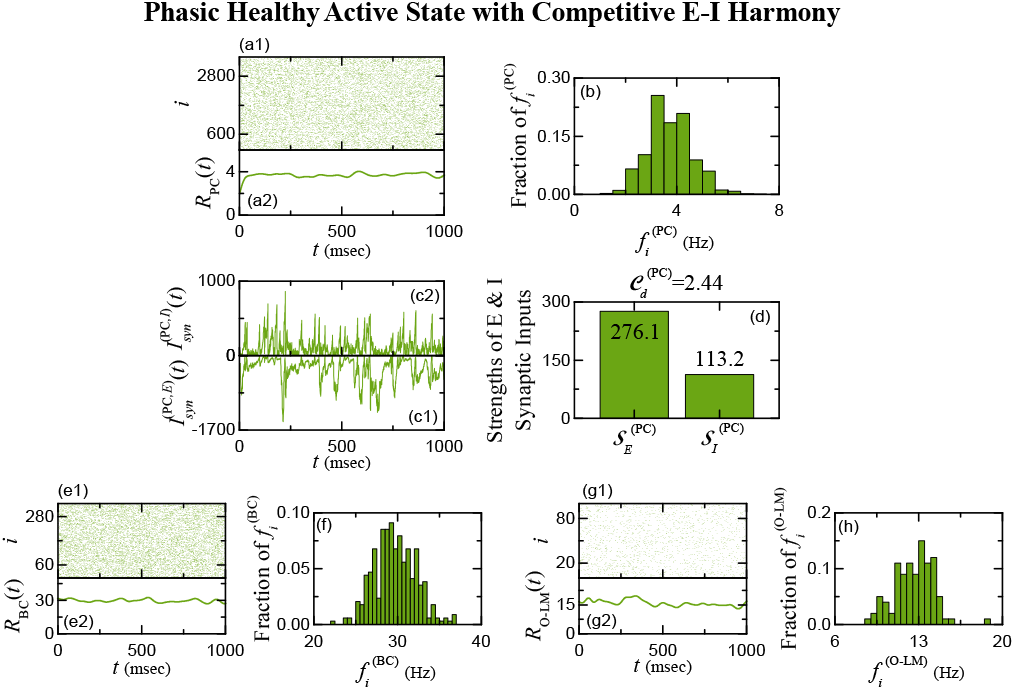
Output-side firing activities (effect) and input-side synaptic quantities (cause) for the phasic healthy active state with competitive E-I harmony (*f*_EC_ = 8 Hz and *f*_CA3_ = 3 Hz). Output-side population and individual firing activities of PCs: (a1) Raster plot of spikes and (a2) IPSR *R*_PC_(*t*) of PCs. (b) Histogram of individual MFRs 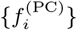 of PCs. Input-side synaptic quantities of PCs: Time series of population-averaged (c1) excitatory (E) 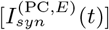 and (c2) inhibitory (I) 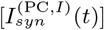 synaptic currents over all PCs. (d) Strengths 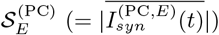 and 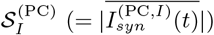 of the population-averaged excitatory and inhibitory synaptic currents. Output-side population and individual firing activities of interneurons: (e1) Raster plot of spikes and (e2) IPSR *R*_BC_(*t*) of BCs. (f) Histogram of individual MFRs 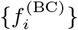 of BCs. (g1) Raster plot of spikes and (g2) IPSR *R*_O*−*LM_(*t*) of O-LM cells. (h) Histogram of individual MFRs 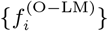 of O-LM cells.

As in the case of tonic resting state, spikes in the raster plots of spikes of the PCs, BCs, and O-LM cells are randomly scattered; consequently, their IPSRs, *R*_PC_(*t*), *R*_BC_(*t*), and *R*_O*−*LM_(*t*) are noisy and irregular, fluctuating around their time averaged values, 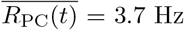, 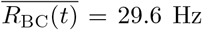, and 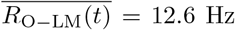. We note that the time-averaged values of IPSRs of PCs, BCs, and O-LM cells are higher than those for the tonic resting state.

Histograms of individual MFRs of PCs, BCs, and O-LM cells are well shown in Figs. 3(b), 3(f), and 3(h), respectively. Their population-averaged MFRs, 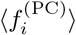, 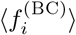, and 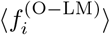, are the same as their above time-averaged IPSRs. Therefore, PCs, BCs, and O-LM cells in the phasic healthy active state exhibit more active firing activities with higher MFRs than those in the tonic resting state.

As input-side quantities (cause) underlying the outputside firing activities (effect) of PCs, time series of population-averaged excitatory and inhibitory synaptic currents, 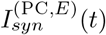 and 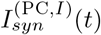, into PCs are shown in Figs. 3(c1) and 3(c2), respectively. Strengths of the excitatory and inhibitory synaptic inputs to PCs, *S*_*E*_ and *S*_*I*_, corresponding to time-averaged magnitudes of 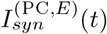 and 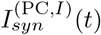, are given in Fig. 3(d); 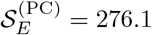 and 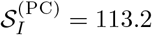. Consequently, the input-side competition degree, 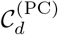, is 2.44 because 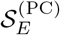 is 2.44 times stronger than 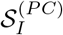, in contrast to the tonic resting state with E-I balance. In this case, both excitatory and inhibitory synaptic inputs to PCs not only compete but also cooperate harmoniously, giving rise to a state of competitive E-I harmony. This corresponds to the phasic healthy active state with 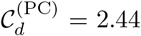, in which the CA1 exhibits stable normal physiological function. In contrast, in the tonic resting state with 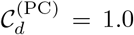, the CA1 remains quiescent with E–I balance, lacking active physiological function.

### B. Breakdown of E-I Harmony via PC Axonal Sprouting: Successive Transitions from HP to IIP to IP

Hyperexcitation and hypoinhibition have long been considered fundamental mechanisms underlying epileptogenesis [6–16]. Two major structural alterations contributing to these mechanisms and thereby leading to the breakdown of E-I harmony are (i) axonal sprouting of PCs, which enhances recurrent excitatory connectivity [33–40], and (ii) loss of inhibitory interneurons, specifically BCs and O-LM cells [40–45].

We first focus on the structural alteration induced by axonal sprouting of PCs. Hippocampal sclerosis is associated with recurrent seizures and progressive loss of PCs. In response to this neuronal loss, the hippocampal network undergoes synaptic reorganization through structural plasticity. Consequently, axonal sprouting enhances recurrent excitatory connectivity among surviving PCs [33–40]. To model this structural alteration, we increase the recurrent excitatory connection probability 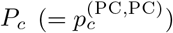 from its normal value of 0.01 and investigate the emergence of pathological states through successive transitions from the healthy phase (HP) to the interictal pathological phase (IIP) and finally to the ictal pathological phase (IP).

Clinically, the interictal, ictal, and postictal phases refer to distinct temporal periods between, during, and after epileptic seizures, respectively [5]. Such temporal evolution of seizure activity has also been investigated in computational studies [65]. In contrast, in our computational study, HP, IIP, and IP represent distinct dynamical states that emerge as the structural pathological parameter *P*_*c*_ is varied, rather than successive temporal stages of seizure activity. For a given value of *P*_*c*_, the CA1 network exhibits a single dynamical state, classified as HP, IIP, or IP according to its firing characteristics. A similar parameter-dependent approach was also employed in a previous computational study, which demonstrated the emergence of synchronized interictal activity under high recurrent axonal sprouting [57].

Figure 4 illustrates the emergence of pathological states through successive transitions from HP (green) to IIP (yellow), and finally to IP (red). In HP [*P*_*c*_ = 0.10, Fig. 4(a)], PC spikes are randomly scattered in the raster plot, showing no apparent population synchronization. Accordingly, the IPSR *R*_PC_(*t*) exhibits noisy and irregular fluctuations, with a low time-averaged value of 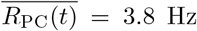. As *P*_*c*_ increases, mutual excitation among PCs becomes stronger. When *P*_*c*_ reaches the first critical threshold 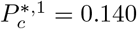, an abrupt transition from HP to IIP occurs due to the constructive effect of moderate hyperexcitation, which arises from moderately increased mutual excitation among PCs and promotes population synchronization.

**FIG. 4:**
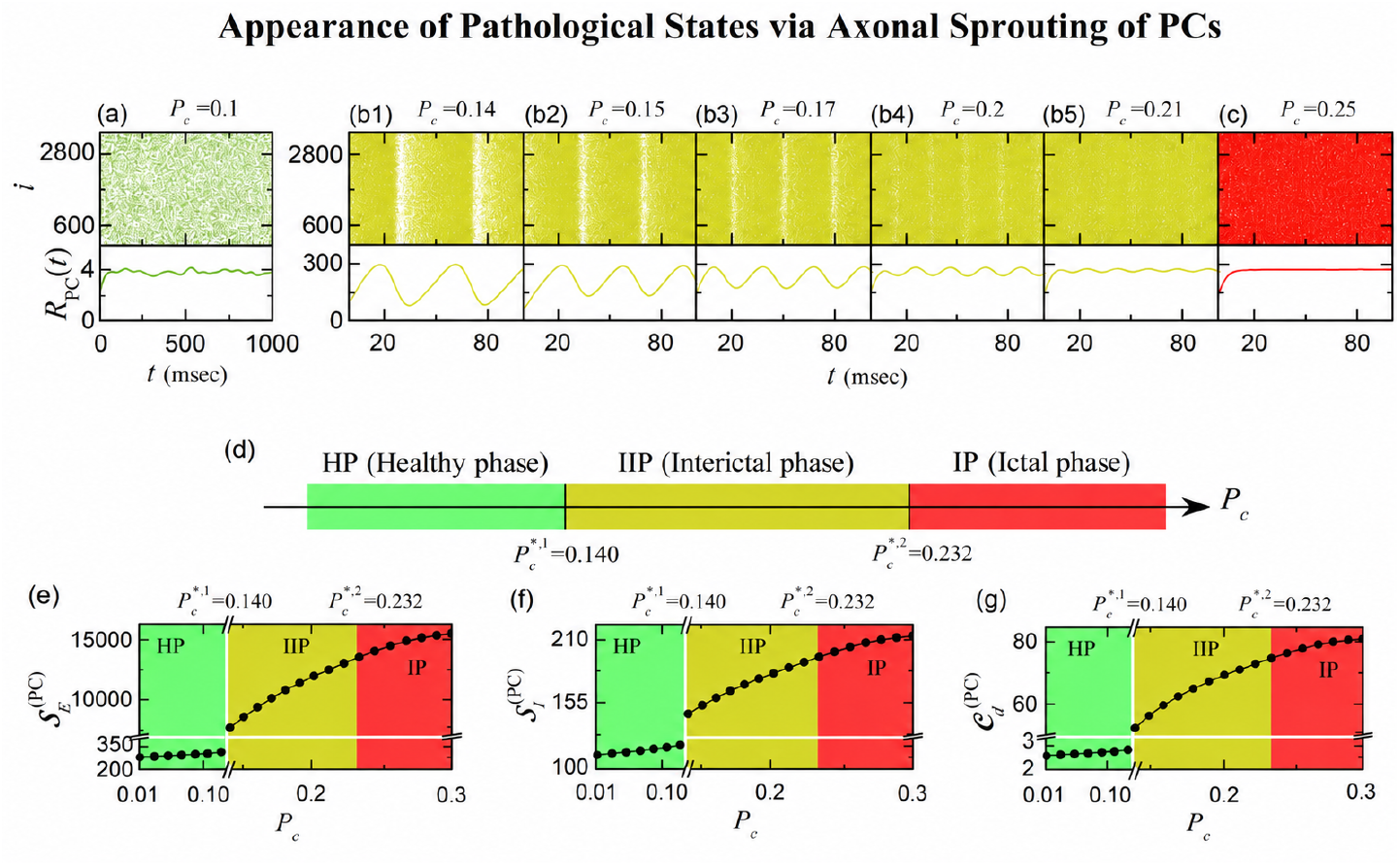
Appearance of pathological states via axonal sprouting of PCs. Output-side population firing activities of PCs (effect): Raster plots of spikes and IPSRs *R*_PC_(*t*) of PCs for the (a) HP (green: *P*_*c*_ = 0.1), (b1)-(b5) IIP (yellow: *P*_*c*_ = 0.14, 0.15, 0.17, 0.2, and 0.21), and (c) IP (red: *P*_*c*_ = 0.25). (d) Bar diagram for the successive transitions from HP to IIP and then to IP. Green, yellow, and red colors represent HP, IIP, and IP, respectively. Input-side synaptic quantities and E-I competition degree of PCs (cause): Plots of strengths (e) 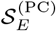 and (f) 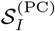 of the population-averaged excitatory and inhibitory synaptic currents over all PCs versus *P*_*c*_. (g) Plot of competition degree 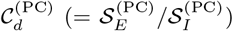 versus *P*_*c*_. For clear presentation, we use a horizontal break for *P*_*c*_ ∈ (0.139, 0.14), a vertical break for 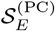 ∈ (400, 8500), and a break for 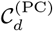 ∈ (3.3, 58).

The emergence of synchronized interictal activity through enhanced recurrent excitation is also consistent with the previous computational findings of Ratnadurai-Giridharan et al. [57] (see their Fig. 8). In IIP [*P*_*c*_ = 0.14, Fig. 4(b1)], synchronized spike bands are clearly visible in the raster plot, and the IPSR *R*_PC_(*t*) exhibits regular, large-amplitude oscillations with a markedly increased time-averaged value of 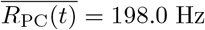. This IIP may reflect features of the clinical interictal phase in epilepsy patients, during which spontaneous, large-amplitude interictal spikes can be observed in EEG recordings in the absence of clinically apparent seizures [57].

With further increases in *P*_*c*_ during IIP [Figs. 4(b2)-4(b5)], the synchronized spike bands in the raster plot gradually approach and merge with one another, leading to a progressive decrease in the degree of population synchronization. This weakening of population synchronization is reflected in the IPSR *R*_PC_(*t*), whose oscillation amplitude gradually decreases. This behavior arises from the destructive role of excessive mutual excitation in population synchronization. Ultimately, when *P*_*c*_ reaches the second critical threshold 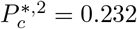, a gradual, continuous transition from synchronized IIP to desynchronized IP is completed through the progressive loss of population synchronization caused by spike-band merging.

Interestingly, this transition from an ordered synchronized IIP to a disordered IP through progressive spike-band merging bears a phenomenological resemblance to the resonance-overlap mechanism in Hamiltonian dynamical systems, where overlapping resonances lead to the breakup of invariant circles and the formation of chaotic regions [107]. Both phenomena exhibit an order-to-disorder transition associated with the overlap of neighboring spike bands or resonances, respectively.

In the desynchronized IP [*P*_*c*_ = 0.25, Fig. 4(c)], PC spikes are randomly scattered in the raster plot, similar to those in HP, showing no apparent population synchronization. However, in sharp contrast to HP, the IPSR *R*_PC_(*t*) exhibits noisy and irregular fluctuations with a very high time-averaged value of 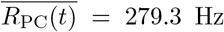. This desynchronized IP state, characterized by irregular and hyperexcitable PC firing, may reflect features of the abnormal neuronal hyperexcitability associated with the clinical ictal phase of epilepsy, during which epileptic seizures occur. The successive transitions from HP (green) to IIP (yellow) and finally to IP (red) are well summarized in the bar diagram in Fig. 4(d).

To uncover the underlying mechanism governing the firing activities of PCs, we next examine the strengths of excitatory and inhibitory synaptic inputs to PCs, 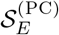 and 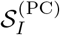, defined in Eq. (6). We then analyze their ratio, the competition degree 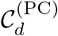 defined in Eq. (7), to quantify E-I harmony and its breakdown across the successive transitions from HP to IIP and then to IP. We note that 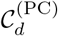 serves as the input-side indicator (cause) underlying the output-side firing activities (effect) of PCs.

While studies typically focus on output-side firing activities (effect), here we emphasize the input-side synaptic quantities (cause)—excitatory and inhibitory synaptic drives— because they directly shape the observed population and individual firing behaviors. By analyzing 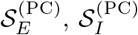, and their ratio (i.e. input-side indicator), the competition degree 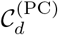, we aim to uncover the mechanistic causes underlying the changes in output-side population and individual firing activities.

Figures 4(e) and 4(f) show variations of 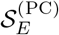 and 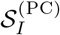 across the three states, HP (green), IIP (yellow), and IP (red), respectively. We first examine 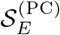. In the HP regime (*P*_*c*_ *<* 0.14), the excitatory synaptic input to PCs increases slowly (e.g., 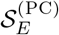= 305.70 at *P*_*c*_ = 0.1), reflecting stable, healthy network activity. At the first threshold 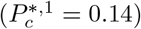, the transition from HP to IIP is accompanied by an abrupt jump in 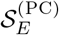 to 9056.24, resulting from intensified recurrent excitation due to PC axonal sprouting. This discontinuous increase marks the onset of a pathological, non-seizure state. Beyond the first threshold, 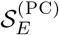 continues to increase gradually through IIP and IP, with the latter characterized by seizure-like activity associated with excessive excitatory drive.

We next examine the variation in 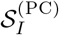. The inhibitory synaptic input to PCs also increases overall with *P*_*c*_, as interneurons are increasingly driven by the enhanced excitatory activity of PCs. In the HP regime, 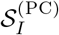 = 117.35 at *P*_*c*_ = 0.1. At the first threshold (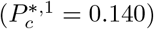= 0.140), corresponding to the transition from HP to IIP, 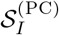 exhibits a small abrupt jump to 150.96, reflecting increased activity of interneurons. Beyond this threshold, 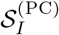 continues to increase gradually, but at a much slower rate than 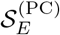. Consequently, throughout IIP and IP, 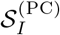 remains much smaller than 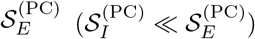, indicating that inhibitory drive cannot keep pace with the rapidly strengthening excitation. Figure 4(g) shows the competition degree 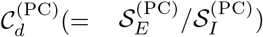, which quantifies the competitive interplay between excitatory and inhibitory synaptic inputs to PCs. In the HP regime, 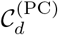 = 2.61 at *P*_*c*_ = 0.1, indicating a state of competitive E–I harmony. In this healthy active state, excitation and inhibition are not perfectly balanced but instead interact cooperatively and competitively to sustain functional network activity. In contrast, perfect E–I balance 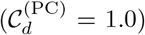 corresponds to the tonic resting state, without physiological functional activities. At the first threshold 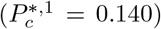, corresponding to the transition from HP to IIP, 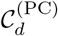 undergoes an abrupt jump to 59.99, arising from the disproportionate increase in excitatory synaptic strength relative to the modest increase in inhibitory synaptic strength. Beyond this threshold, 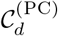 increases gradually with *P*_*c*_ throughout IIP and IP, indicating the breakdown of competitive E–I harmony and the emergence of excitation-dominated disharmony characteristic of pathological states.

Having examined the input-side E–I competition in Fig. 4, we now turn to the output-side firing activities of PCs (effect). Figure 5 presents the changes in PC firing activities across the successive transitions from HP to IIP and then to IP, providing an output-side characterization of the physiological and pathological states. We first characterize the population firing activities of PCs using two measures: the time-averaged IPSR 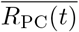 (the over-line denotes time averaging) and the amplitude measure 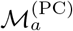, which quantifies the degree of population synchronization [108, 109]. Each data point was obtained from a 10,000-msec numerical simulation.

**FIG. 5:**
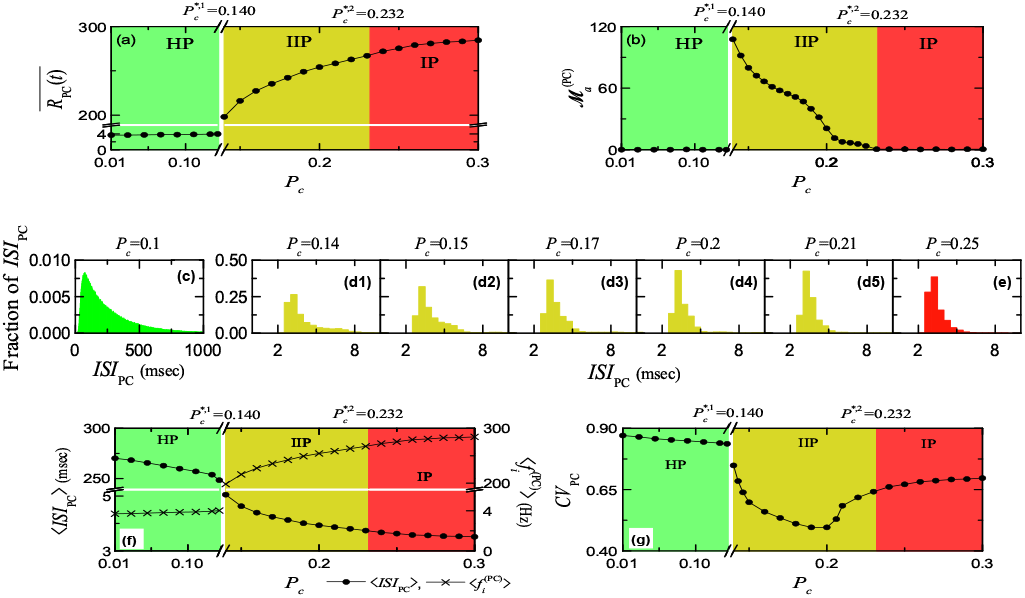
Population and individual firing activities of PCs in the healthy and pathological states via axonal sprouting of PCs. Output-side population firing activities of PCs: (a) Plot of time-averaged IPSR 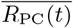 versus *P*_*c*_. For clear presentation, we use a horizontal break with *P*_*c*_ ∈ (0.139, 0.14) and a vertical break for 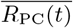 ∈ (6, 190). (b) Plot of amplitude measure 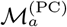 versus *P*_*c*_. Output-side individual firing activities of PCs: Population-averaged ISI (interspike interval) histograms for (c) HP (green: *P*_*c*_ = 0.1), (d1)-(d5) IIP (yellow: *P*_*c*_ = 0.14, 0.15, 0.17, 0.2, and 0.21), and (e) IP (red: *P*_*c*_ = 0.25). (f) Plots of average ISI ⟨ ISI_PC_ ⟩ (circles) and population-averaged MFR (mean firing rate) 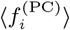 (= 1*/*⟨ ISI_PC_ ⟩) (crosses) versus *P*_*c*_. (g) Plot of *CV*_PC_ (coefficient of variation) versus *P*_*c*_. For clear presentation in (f) and (g), we use a horizontal break with *P*_*c*_ ∈ (0.139, 0.14) and also use vertical breaks for ⟨ ISI_PC_ ⟩ ∈ (5.2, 240) and 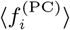 ∈ (6, 190) in (f).

Figure 5(a) shows the variation of the time-averaged IPSR 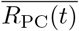 across the three states: HP, IIP, and IP. In the HP regime, 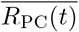 increases very slowly with *P*_*c*_, starting from 3.7 Hz at *P*_*c*_ = 0.01. At the first threshold 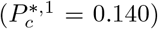, however, 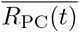 exhibits an abrupt jump to 198.0 Hz, marking the transition from HP to IIP. Beyond this threshold, 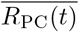 continues to increase gradually with *P*_*c*_ throughout IIP and IP. The amplitude of the IPSR *R*_PC_(*t*) reflects the degree of population synchronization. To quantify this, we employ the amplitude measure 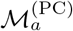, defined as the time-averaged amplitude of *R*_PC_(*t*) [108, 109]:

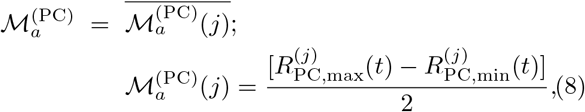

where the overline denotes time averaging, 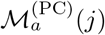 is the amplitude measure in the *j*th global cycle of *R*_PC_(*t*) (corresponding to the *j*th spike band in the raster plot). Here, 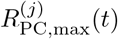 and 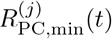 are the maximum and minimum values of *R*_PC_(*t*) during the *j*th global cycle, respectively. Larger values of 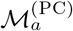 indicate stronger oscillatory modulation of *R*_PC_(*t*) and, consequently, a higher degree of population synchronization.

Variation of the synchronization degree, quantified by 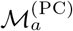, is shown across the three states (HP, IIP, and IP) in Fig. 5(b). As shown in Fig. 4, no population synchronization occurs in either HP or IP, because spikes in their raster plots are randomly scattered without forming spike bands (i.e., their IPSRs *R*_PC_(*t*) exhibit noisy, irregular fluctuations without regular oscillations). Consequently, 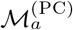 = 0 in both HP and IP. In contrast, during IIP, synchronized spike bands appear successively, giving rise to regular oscillation of the IPSR *R*_PC_(*t*). At the 1st threshold, 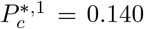, an abrupt jump in 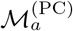 to its maximum value (107.6) occurs due to the constructive effect of moderate increase in mutual excitation, which promotes the emergence of synchronized spike bands. However, with further increases in *P*_*c*_ during IIP,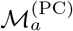 gradually decreases because of the destructive effect of excessive mutual excitation, which weakens population synchronization. Eventually, 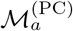 falls to nearly zero at the 2nd threshold, 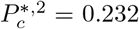, marking the transition to IP.

In the IIP regime, we also examine the population synchronization frequency 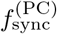, corresponding to the oscillation frequency of the IPSR *R*_PC_(*t*). It is calculated as the inverse of the time-averaged oscillation period of *R*_PC_(*t*), obtained from successive global cycles during the 10,000-msec numerical simulation.

The population synchronization frequency 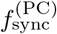 increases monotonically with increasing *P*_*c*_. For example, 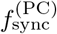 is 22.8 Hz at *P*_*c*_ = 0.14 and 57.1 Hz at *P*_*c*_ = 0.21. In contrast, the synchronization degree, quantified by 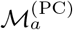, gradually decreases with increasing *P*_*c*_. Thus, as mutual excitation among PCs increases during IIP, the population oscillations become faster but progressively weaker in amplitude.

In addition to population firing activity, we also analyze individual firing activity of PCs across HP, IIP, and IP. For this purpose, we use two measures: the mean interspike interval (ISI), ⟨ *ISI*_PC_ ⟩, and the coefficient of variation *CV*_PC_, which quantifies the temporal irregularity of individual firing. Here, *CV*_PC_ is defined as *CV*_PC_ = *SD*_PC_*/ ⟨ ISI*_PC_ ⟩, where *SD*_PC_ denotes the standard deviation of the ISIs, measuring their spread around the mean ⟨ *ISI*_PC_ ⟩. For each PC, ISIs were extracted from a 10,000-ms numerical simulation.

Figures 5(c)-5(g) summarize these results: panels (c)–(e) show population-averaged ISI histograms for HP (green), IIP (yellow), and IP (red), while panels (f)–(g) display the corresponding variations of ⟨ *ISI*_PC_ ⟩ and *CV*_PC_. Along with ⟨ *ISI*_PC_ ⟩ (circles), the population-averaged MFRs 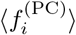 (crosses), given by the reciprocal of ⟨ ISI_PC_ ⟩, are also shown in Fig. 5(f).

In HP (green), ⟨ ISI_PC_ ⟩ decreases gradually with increasing *P*_*c*_ (e.g., at *P*_*c*_ = 0.1, ⟨ ISI_PC_ ⟩ = 260.5 msec), while 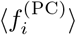 rises slowly (e.g., 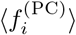 = 3.8 Hz for *P*_*c*_ = 0.1). However, at the 1st threshold (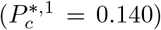 = 0.140) for the HP–IIP transition, a sudden jump occurs: ⟨ *ISI*_PC_ ⟩ drops to 5.05 msec, while 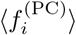 increases sharply to 198.0 Hz. Thereafter, during IIP and IP, ⟨ ISI_PC_ ⟩ continues to decrease and 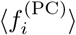 continues to increase, leading to hyperexcitable firing in IP (e.g., at *P*_*c*_ = 0.25, 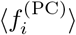 = 279.3 Hz, corresponding to ⟨ ISI_PC_ ⟩ = 3.58 msec).

We next consider the coefficient of variation, *CV*_PC_ [Fig. 5(g)]. In HP, PCs exhibit highly irregular firing, as reflected by large *CV*_PC_ values (e.g., *CV*_PC_ = 0.85 at *P*_*c*_ = 0.1). With increasing *P*_*c*_, *CV*_PC_ decreases gradually, followed by a sudden drop to *CV*_PC_ = 0.75 at the 1st threshold of 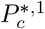 = 0.140. Within IIP, *CV*_PC_ continues to fall rapidly and reaches a minimum of 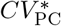= 0.492 at 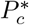 = 0.194. For larger 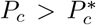, spanning the late IIP and IP, *CV*_PC_ rises again, forming a well-defined U-shaped profile and indicating a progressive increase in firing irregularity.

Comparing the population and individual firing activities of PCs in Fig. 5 reveals an interesting contrast between network-level synchronization and single-neuron firing regularity. In IIP, population synchronization steadily decreases with increasing *P*_*c*_, whereas the irregularity of individual neuron firing (*CV*_*PC*_⟩ decreases specifically in the range 0.14 *< P*_*c*_ *<* 0.194. At first glance, this might seem counterintuitive, as one could expect that more regular individual firing would enhance collective synchrony. However, population synchronization depends not only on the regularity of each neuron’s spikes but also on the temporal alignment of spikes across the network. In this intermediate regime, individual neurons fire more regularly, yet the network lacks sufficient coordination to align spikes, leading to reduced population synchronization. This highlights the importance of distinguishing single-neuron firing activity from network-level coordination when interpreting trends in the coefficient of variation (*CV*_*PC*_) and the synchronization degree 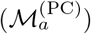.

Another distinctive feature of population synchronization in CA1 is the multiple spiking of individual PCs within each synchronized band, corresponding to a global cycle of the IPSR. This dense population synchronization contrasts with sparse synchronization, where only a fraction of neurons fire during each global cycle due to spike skipping [101–105]. Accordingly, in CA1, the population synchronization frequency is lower than the population-averaged MFR of individual PCs, 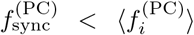, whereas the opposite relation, *f*_sync_ *> ⟨ f*_*i* ⟩_, holds for sparse synchronization.

### C. Breakdown of E-I Harmony via Interneuron Loss: Successive Transitions from HP to IIP to IP

In this subsection, we focus on the structural pathological alteration caused by the loss of inhibitory interneurons, which leads to the breakdown of E-I harmony through hypoinhibition. Hippocampal sclerosis is associated with recurrent seizures and a progressive loss of interneurons. To capture this effect, we reduce the numbers of BCs and O-LM cells from their original values, 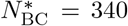 and 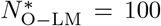. The surviving populations are scaled by a common survival fraction

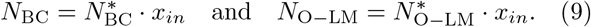

Progressively decreasing *x*_*in*_ (fraction of surviving interneurons) from 1 represents increasing interneuron loss. For a fixed recurrent connection probability *P*_*c*_ = 0.1 between PCs, we examine how this structural pathological change (i.e., loss of interneurons) leads to the breakdown of E-I harmony and successive transitions from HP (healthy phase) to IIP (interictal phase) and finally to IP (ictal phase).

Figures 6(a)-6(c) show raster plots of PC spikes and the corresponding IPSRs *R*_PC_(*t*) for representative values of the surviving interneuron fraction *x*_in_. In HP [green; e.g., Fig. 6(a) for *x*_in_ = 0.8], PC spikes are randomly scattered in the raster plot, resulting in a noisy and irregular IPSR *R*_PC_(*t*) with a low time-averaged value, 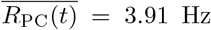. As shown in Fig. 6(d), when *x*_in_ decreases to the first threshold 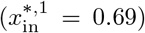, the network undergoes an abrupt transition from HP to IIP. In IIP [yellow; Figs. 6(b1)–6(b5)], synchronized spike bands emerge in the raster plots, accompanied by regular oscillations in *R*_PC_(*t*). With further reduction of *x*_in_, neighboring spike bands become progressively closer and overlap, leading to a gradual decrease in the degree of population synchronization.

**FIG. 6:**
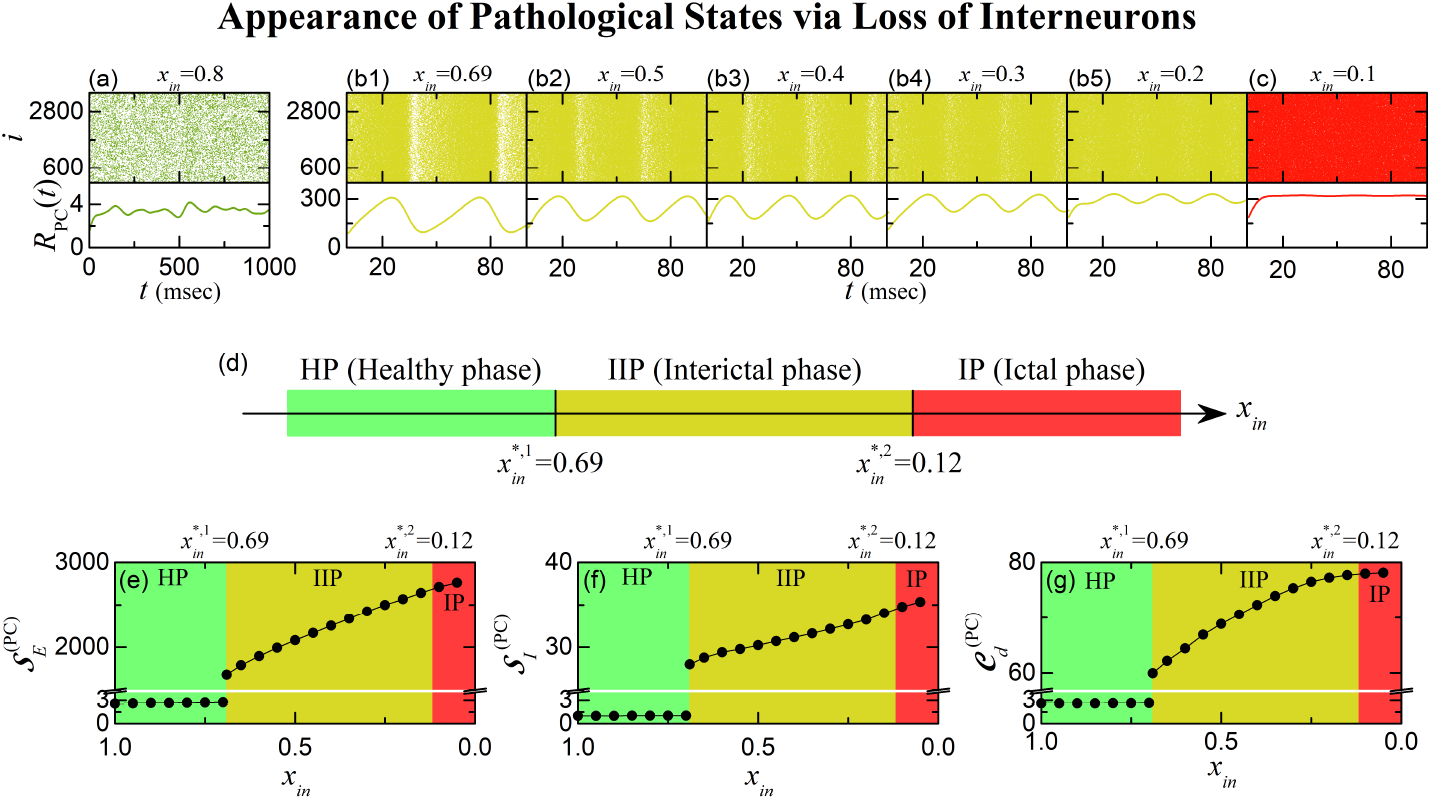
Appearance of pathological states via loss of interneurons (BCs and O-LM cells) for *P*_*c*_ = 0.1. Output-side population firing activities of PCs (effect): Raster plots of spikes and IPSRs *R*_PC_(*t*) of PCs for the (a) HP (green: *x*_*in*_ = 0.8), (b1)-(b5) IIP (yellow: *x*_*in*_ = 0.69, 0.5, 0.4, 0.3, and 0.2), and (c) IP (red: *x*_*in*_ = 0.1). (d) Bar diagram for the successive transitions from HP to IIP and then to IP. Green, yellow, and red colors represent HP, IIP, and IP, respectively. Input-side synaptic quantities and E–I competition degree of PCs (cause): Plots of strengths (e) 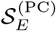 and (f) 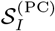 of the population-averaged excitatory and inhibitory synaptic currents into PCs versus *x*_*in*_. (g) Plot of competition degree 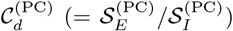 versus *x*_*in*_. For clear presentation, we use vertical breaks for 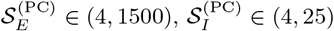, and a break for 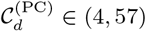.

Ultimately, at the second threshold [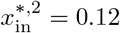, shown in Fig. 6(d)], the synchronized spike bands become completely merged, and the network enters desynchronized IP [red; e.g., see Fig. 6(c) for *x*_in_ = 0.1]. In IP, individual PC spikes become highly irregular and scattered in the raster plot, resulting in no population synchronization. Consequently, the IPSR *R*_PC_(*t*) exhibits irregular fluctuations with a very high time-averaged value (e.g., 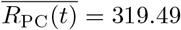 Hz at *x*_in_ = 0.1).

To uncover the underlying mechanism governing the emergence of pathological states caused by interneuron loss, we next examine the strengths of excitatory and inhibitory synaptic inputs to PCs, 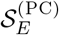 and 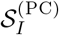, and their ratio, the competition degree 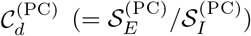. As in the case of PC axonal sprouting, 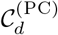 serves as the input-side indicator characterizing E-I harmony and its breakdown across the successive transitions from HP to IIP and then to IP.

Figures 6(e)-6(g) quantitatively show how interneuron loss alters excitatory and inhibitory synaptic inputs to PCs and their competitive interplay, quantified by the competition degree 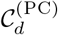. In Fig. 6(e), the strength 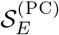 of excitatory synaptic input to PCs shows a gradual increase with decreasing *x*_in_, reflecting disinhibition of the PC population due to hypoinhibition (i.e., reduced interneuron control). Somewhat counterintuitively, the strength 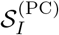 of inhibitory synaptic input to PCs in Fig. 6(f) also exhibits a small overall increase, despite the loss of interneurons. This paradoxical rise arises because surviving interneurons receive stronger excitatory drive from the increasingly active PCs, thereby partially compensating for their reduced numbers. Compared with PC axonal sprouting, both 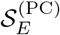 and 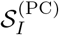 remain substantially lower in magnitude.

Finally, Fig. 6(g) shows the competition degree ^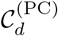^, quantifying competitive interplay between excitatory and inhibitory synaptic inputs to PCs. At the HP–IIP transition 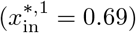, 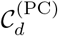 exhibits an abrupt jump from its healthy value (e.g., ^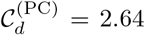^ at *x*_*in*_ = 0.8) to a much larger one (59.96), signifying the onset of a pathological state. Thereafter, during IIP and IP, ^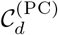^ gradually increases with decreasing *x*_in_, indicating progressive breakdown of competitive E-I harmony. These changes in input-side synaptic quantities (cause) such as 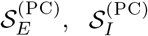, and 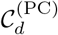 provide the mechanistic basis for the changes in output-side population and individual PC firing activities (effect) shown in Fig. 7.

**FIG. 7:**
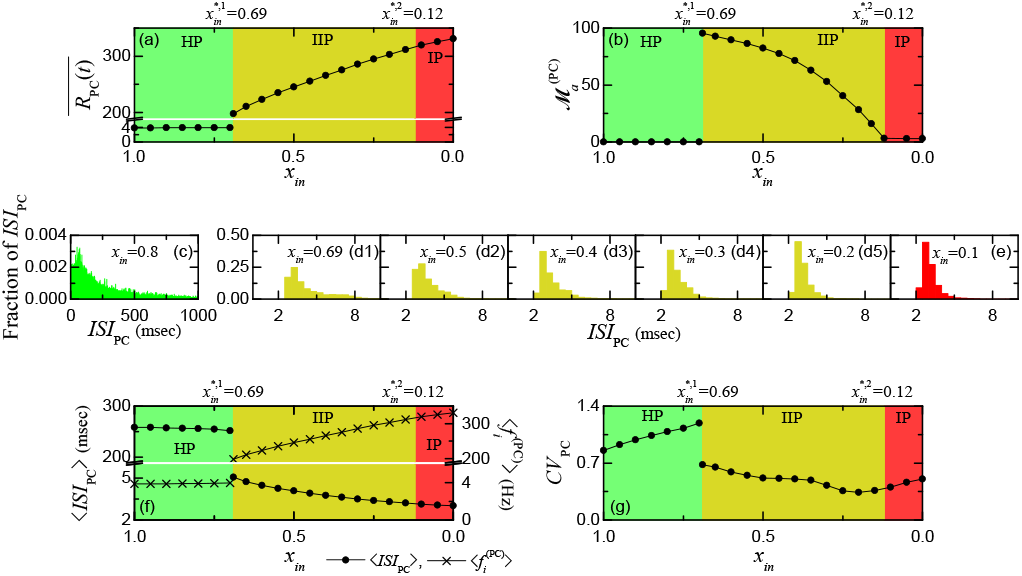
Population and individual firing activities of PCs in the healthy and pathological states via loss of interneurons (BCs and O-LM cells) for *P*_*c*_ = 0.1. Output-side population firing activities of PCs: (a) Plot of time-averaged IPSR 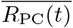 versus *x*_*in*_. For clear presentation, we use a vertical break for 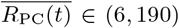. (b) Plot of amplitude measure 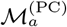 versus *x*_*in*_. Output-side individual firing activities of PCs: Population-averaged ISI (interspike interval) histograms for (c) HP (green: *x*_*in*_ = 0.8), (d1)-(d5) IIP (yellow: *x*_*in*_ = 0.69, 0.5, 0.4, 0.3, and 0.2), and (e) IP (red: *x*_*in*_ = 0.1). (f) Plot of average ISI ⟨*ISI*_*PC*_⟩ (circles) and population-averaged MFR (mean firing rate) 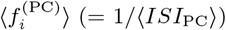 (crosses) versus *x*_*in*_. For clear presentation, we use vertical breaks for ⟨*ISI*_*PC*_⟩ ∈ (6, 190) and 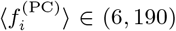. (g) Plot of *CV*_PC_ (coefficient of variation) versus *x*_*in*_.

We now examine the output-side consequences of the input-side E-I competition associated with interneuron loss. Figure 7 shows how the population and individual firing activities of PCs change across the successive transitions from HP to IIP and then to IP. We first consider the population firing activities of PCs. Figs. 7(a) and 7(b) show the variations of the time-averaged IPSR 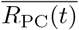 and the amplitude measure 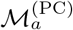, respectively, as functions of the surviving interneuron fraction *x*_in_. During HP (green), 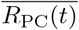 (3.9 Hz) increases very slowly. At the first threshold 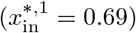 for the transition to IIP, an abrupt jump to 198.02 Hz occurs. There-after, during IIP (yellow) and IP (red), 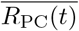 gradually increases with decreasing *x*_in_. In IP, the time-averaged IPSR 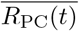 becomes very high (e.g., for *x*_in_ = 0.1, 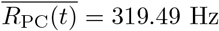).

We also consider the amplitude measure ^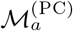^, quantifying the degree of population synchronization. In HP and IP, ^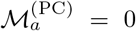^ because population synchronization is absent. When *x*_in_ reaches the first threshold 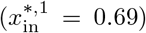 for the transition to IIP, population synchronization emerges abruptly, with ^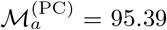^, due to the constructive effect of moderate hypoinhibition caused by moderate interneuron loss. As *x*_in_ is further decreased from 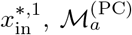 decreases monotonically to nearly zero at the second threshold 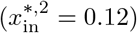 for the transition to IP, reflecting the destructive effect of excessive hypoinhibition caused by severe interneuron loss. Consequently, for 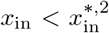, the desynchronized IP appears.

We then turn to the population synchronization frequency 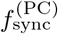, corresponding to the oscillation frequency of the IPSR *R*_PC_(*t*) of the PCs. As *x*_in_ decreases in the IIP regime, 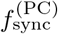 increases monotonically (e.g., from 20.5 Hz at *x*_in_ = 0.69 to 34.6 Hz at *x*_in_ = 0.20). In contrast, the synchronization degree, quantified by ^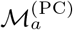^, gradually decreases. Thus, as interneuron loss progresses during IIP, the population oscillations become faster, while the degree of population synchronization progressively decreases. As in the PC axonal sprouting case, individual PCs exhibit multiple spiking within each synchronized band, resulting in dense population synchronization. Accordingly, the population synchronization frequency is lower than the population-averaged MFR of individual PCs, 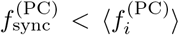, whereas the opposite relation, *f*_sync_ *> ⟨ f*_*i*_ ⟩, holds for sparse population synchronization arising from spike skipping. The numerical values of 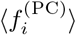 are presented below in the analysis of individual PC firing activities.

Next, we consider the individual firing activities of the PCs. Panels (c)-(e) present the ISI histograms for representative values of *x*_*in*_ in HP (green), IIP (yellow), and IP (red). We analyze the individual firing behavior of PCs using two measures: the average ISI ⟨ *ISI*_PC_ ⟩, and the coefficient of variation *CV*_PC_ (quantifying the irregularity degree of individual firings). The variations of ⟨ *ISI*_PC_ ⟩ and *CV*_PC_ are shown in panels (f) and (g), respectively. Together with ⟨*ISI*_*PC*_⟩ (circles), the population-averaged 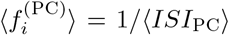 (crosses) are also shown in panel (f).

In HP, ⟨*ISI*_*PC*_⟩ decreases slowly with decreasing *x*_*in*_ (e.g., at *x*_*in*_ = 0.8, ⟨*ISI*_*PC*_⟩ = 255.56 msec), while 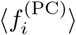 rises slowly (e.g., at *x*_*in*_ = 0.8, 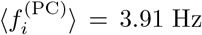. At the 1st threshold of 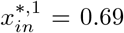 for the transition to IIP, an abrupt jump occurs: ⟨*ISI*_*PC*_⟩ drops to 5.05 msec, while 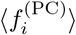 rises sharply to 198.02 Hz. Thereafter, during IIP and IP, ⟨*ISI*_PC_ ⟩ continues to decrease and 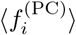 continues to increase, resulting in hyperexcitable firings in IP (e.g., at *x*_*in*_ = 0.1, 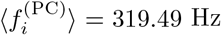, corresponding to ⟨*ISI*_*PC*_⟩ = 3.13 msec).

The behavior of *CV*_PC_ is particularly noteworthy: in contrast to the axonal sprouting case (where *CV*_PC_ decreases in HP), here *CV*_PC_ increases already during HP, and exceeds 1 (implying more irregular than a Poisson-like process). At the 1st threshold of 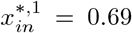 for the transition to IIP, *CV*_PC_ drops suddenly. In the subsequent IIP and IP, *CV*_PC_ exhibits a U-shaped (bowl-shaped) profile—initially decreasing but rising again in deeper pathological states—indicating an initial reduction in firing irregularity followed by a subsequent increase.

Together, these results show that interneuron loss induces both increased firing rates and changes in the temporal irregularity of individual PCs. These firing patterns are largely similar to those in the sprouting case—showing a U-shaped profile of the *CV*_PC_ during IIP and IP—but they differ in HP, where the *CV*_PC_ increases under interneuron loss, in contrast to its decrease in the sprouting case.

We also compare the final epileptic state (IP generated by interneuron loss) with IP generated by the axonal sprouting of PCs. Both PC sprouting and interneuron loss can drive the network into pathological states (IP), but the resulting IP states differ in severity. For example, at *P*_*c*_ = 0.25 (IP induced by PC sprouting), the competition degree is 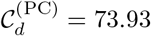 and the population-averaged MFR is 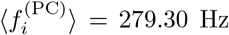, whereas for interneuron loss at *x*_*in*_ = 0.1 (IP), 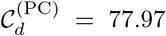 and 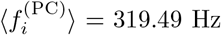. This indicates that IP induced by interneuron loss exhibits stronger pathological hyperexcitability. The apparent paradox—larger 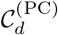 despite smaller strengths, 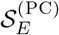 and 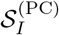, of the excitatory and inhibitory synaptic inputs to PCs in interneuron loss—arises because ^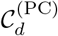^ reflects the relative disharmony between excitation and inhibition. Even though both 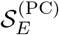 and 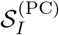 are smaller, their ratio becomes larger, indicating more severe E-I disharmony, consistent with the observed stronger pathological hyperexcitability of PCs.

### D. Unified State Diagram of Epileptogenesis in the P_c_ −*x*_*in*_ Plane

Having analyzed separately the emergence of pathological states via axonal sprouting of PCs (Figs. 4-5) in Sec. III B and loss of inhibitory interneurons (Figs. 6-7) in Sec. III C, we now present the overall state diagram in the *P*_*c*_ - *x*_*in*_ plane (see Fig. 8). This state diagram integrates both structural pathological mechanisms into a unified two-dimensional framework. The three regimes— HP (green), IIP (yellow), and IP (red)—occupy distinct regions, with boundaries determined by the input-side competition degree 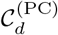 for the HP → IIP transition and by the output-side amplitude measure 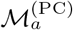 for the IIP → IP transition.

**FIG. 8:**
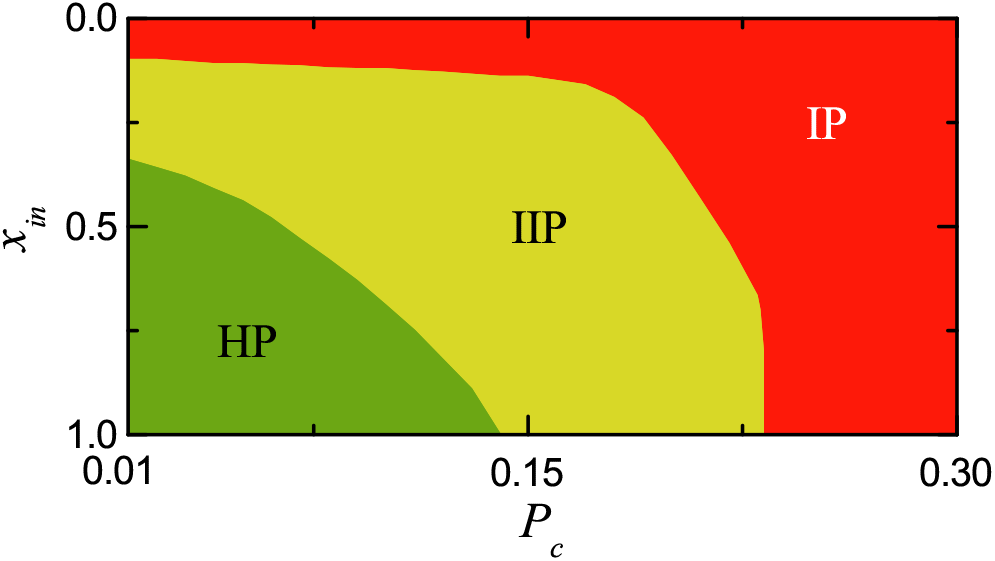
Unified overall state diagram in the *P*_*c*_-*x*_*in*_ plane for the combined effects of axonal sprouting of PCs and loss of interneurons (BCs and O-LM cells). Green, yellow, and red colors denote the healthy phase (HP), interictal pathological phase (IIP), and ictal pathological phase (IP), respectively.

The unified state diagram clearly illustrates that structural changes in the CA1 network, whether by PC axonal sprouting, interneuron loss, or their combination, result in the onset of pathological states. Moreover, PC sprouting and interneuron loss can jointly trigger the abrupt transition from HP to IIP through moderate changes in both variables, while more severe alterations drive the transition from IIP into IP.

Thus, Fig. 8 provides a unified quantitative characterization of the onset and progression of pathological states in the two-dimensional structural parameter space. In particular, the HP → IIP transition marks the onset of pathological states due to the breakdown of E-I harmony, accompanied by a discontinuous, abrupt jump in the competition degree 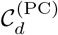, which serves as a potential quantitative biomarker of this transition.

## IV. SUMMARY AND DISCUSSION

We developed a biologically grounded SNN model of hippocampal CA1 to investigate how two structural pathological mechanisms associated with epileptogenesis—axonal sprouting of PCs and loss of inhibitory BCs and O-LM cells—drive the breakdown of E-I harmony. As recurrent excitatory connectivity increases through PC axonal sprouting or inhibitory interneuron populations decline through interneuron loss, the network undergoes sequential transitions from a healthy phase (HP) to an interictal pathological phase (IIP) and finally to an ictal pathological phase (IP).

To quantitatively characterize these transitions, we introduced the competition degree 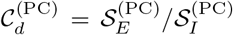, defined as the ratio of the strengths of excitatory to inhibitory synaptic inputs to PCs, as an input-side indicator of competitive E-I harmony. In HP, 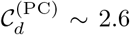, reflecting stable E-I harmony sustained by cooperative interplay between excitation and inhibition. At the HP → IIP transition, 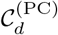 exhibited an abrupt, discontinuous jump to *~*60, followed by a further gradual increase through IIP and IP. Correspondingly, the output-side measure, the population-averaged MFR of PCs, 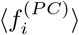, also exhibited an abrupt, discontinuous jump from *~*4 to *~*200 Hz at the HP → IIP transition and increased further thereafter, indicating progressive pathological hyperexcitability. Thus, 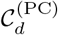 may serve as a potential quantitative biomarker of competitive E-I harmony and its breakdown during epileptogenesis.

Although PC axonal sprouting and interneuron loss alter CA1 circuitry through distinct structural mechanisms, both lead to the breakdown of competitive E-I harmony, as quantified by the input-side indicator 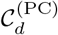, accompanied by increased output-side PC firing activity. Thus, these two distinct pathological routes converge on a common quantitative cause-and-effect framework for the progression from healthy to pathological states.

Experimentally, 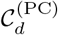 could potentially be estimated in animal models or appropriate experimental preparations using established electrophysiological techniques. Dual or triple patch-clamp recordings in voltage-clamp mode can separately measure excitatory and inhibitory postsynaptic currents [66, 67]. The time-averaged magnitudes of these currents can provide values of 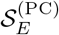 and 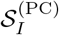, whose ratio gives 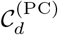. Alternatively, imaging methods such as calcium or voltage-sensitive dye imaging combined with cell-type-specific labeling [68, 69] may provide indirect estimates of excitatory and inhibitory synaptic activity, offering potential avenues toward in vivo studies and eventual clinical applications.

Detection of a discontinuous, abrupt jump in 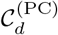 at the HP → IIP transition suggests critical intervention windows where targeted therapies—such as optogenetic modulation of CA1 circuits [110], pharmacological enhancement of inhibition [111], or suppression of pathological pyramidal-cell sprouting [112]—have the potential to restore E-I harmony, thereby returning the pathological network toward the healthy state.

As structural alterations accumulate, excitation increasingly dominates over inhibition, reflecting the progressive breakdown of E-I harmony. In the PC sprouting case, moderately increased mutual excitation among PCs leads to moderate hyperexcitation, whose constructive effect promotes population synchronization and triggers the HP → IIP transition. In the interneuron loss case, moderate hypoinhibition caused by moderate interneuron loss plays a corresponding constructive role, leading to the emergence of IIP. In both cases, the emergence of population synchronization marks the HP → IIP transition. This intermediate pathological regime resembles the interictal state observed clinically, in which spontaneous, large-amplitude interictal spikes are observed in EEG recordings, and may represent an early manifestation of network instability preceding seizures.

By contrast, excessive mutual excitation among PCs in the sprouting case leads to excessive hyperexcitation, while severe interneuron loss causes excessive hypoinhibition. Both mechanisms exert destructive effects on population synchronization: synchronized population rhythms progressively deteriorate and eventually disappear as the network enters IP, characterized by irregular high-rate PC firing and pathological hyperexcitability. This irregular high-rate PC firing activity may reflect features of the abnormal neuronal hyperexcitability associated with the clinical ictal phase of epilepsy. Thus, the degree of structural alteration plays a dual dynamical role: moderate alterations promote population synchronization, leading to the onset of IIP, whereas severe alterations destroy population synchronization and drive the transition to IP.

Previous studies have qualitatively linked E-I imbalance to epilepsy, emphasizing excitatory sprouting and interneuron loss as key drivers. However, most studies lacked a quantitative measure bridging network structural changes to functional state transitions. Our introduction of 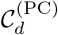 fills this gap by providing a quantitative input-side indicator of competitive E-I harmony and its breakdown, directly linking structural pathological changes to their output-side effects and epileptogenic state transitions. This advances the classical concept of “E-I balance” toward a measurable, quantitative notion of “E-I harmony.”

Our current model focuses on CA1 with external excitatory drive from CA3 and EC modeled as Poisson spike trains. Future work will incorporate additional hippocampal subregions (e.g., CA3 and dentate gyrus), synaptic plasticity mechanisms, and more detailed interneuron modeling. Experimental validation of 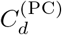 using in vitro and in vivo preparations is essential, and applying 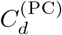 to human hippocampal recordings could help evaluate its potential utility as a clinical biomarker.

Future computational studies will also explore optogenetic strategies to restore E-I harmony by selectively deactivating PCs or activating interneurons, thereby reducing 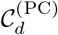 and suppressing pathological hyperexcitability. In the representative IP states examined, interneuron loss produces a more severe pathological state than PC sprouting alone, as reflected by the stronger PC hyperexcitability. This underscores the critical role of inhibitory control in maintaining network stability and supports strategies that enhance interneuron activity as a first step in controlling epileptiform dynamics.

Hippocampal network excitability can be conceptualized in terms of Yin-Yang harmony between inhibition (Yin) and excitation (Yang). The conventional notion of balance may be contrasted with the concept of Yin-Yang harmony, in which contrasting yet complementary elements interact harmoniously without requiring equality. From this perspective, the interaction between excitation and inhibition may be understood as a competitive harmony between Yin (inhibition) and Yang (excitation), rather than as an E-I balance. The competition degree 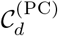 quantitatively characterizes this input-side competitive E-I harmony and its breakdown, while the population-averaged MFR of PCs, 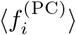, characterizes the resulting output-side firing activity during epileptogenesis.

Finally, our competition-degree-based causal framework, developed to quantify E-I harmony, has broad potential beyond epilepsy (see Fig. 9). In a broader context, excitation (E) and inhibition (I) may be considered as function-promoting (Function) and controlling (Control) influences, respectively. Thus, the competition degree *C*_*d*_ = *E/I* can be viewed more generally as the ratio of Function to Control, quantifying their competitive yet cooperative harmony. This generalized perspective also encompasses our previous studies of the basal ganglia [18, 19], where the direct “Go” pathway promotes action (Function), while the indirect “No-Go” pathway provides inhibitory control (Control). In this sense, the competitive yet cooperative interactions in both the basal ganglia and hippocampal CA1 may be viewed as specific neural manifestations of the broader philosophical concept of Yin-Yang harmony.

**FIG. 9:**
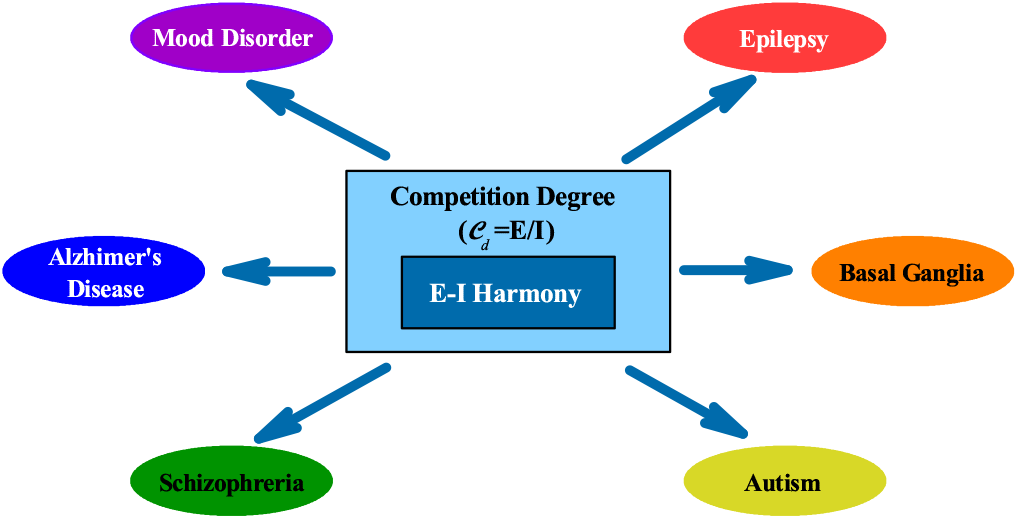
Generalized quantitative framework for competitive yet cooperative E-I harmony and its generalization to Function-Control harmony. The competition degree *C*_*d*_ = *E/I* = Function*/*Control provides an input-side quantitative measure of this harmony and its breakdown, offering a potential common causal framework for understanding diverse neurological and psychiatric disorders, including epilepsy and basal ganglia disorders. Such competitive harmony may be viewed as a neural manifestation of the broader philosophical concept of Yin-Yang harmony.

Interestingly, the healthy-state competition degrees in the basal ganglia and hippocampal CA1 are approximately 2.8 and 2.6, respectively. In both systems, normal physiological function is sustained by competitive yet cooperative harmony, with the function-promoting influence being approximately 2.7 times stronger than the controlling influence. Pathological states emerge when this harmony breaks down through alterations in Function or Control. In the basal ganglia, overactivity of the indirect pathway in Parkinson’s disease decreases *C*_*d*_, whereas its underactivity in Huntington’s disease increases *C*_*d*_. In hippocampal CA1, PC axonal sprouting and interneuron loss increase *C*_*d*_ through hyperexcitation and hypoinhibition, respectively. Thus, distinct pathological mechanisms can be understood within a common causal framework linking input-side competitive disharmony to output-side pathological activity.

E-I disharmony is also implicated in diverse neurological and psychiatric disorders, including autism spectrum disorder, schizophrenia, Alzheimer’s disease, and mood disorders [70–77]. The close numerical values of the healthy-state competition degrees in the basal ganglia (*~* 2.8) and CA1 (*~*2.6) are intriguing, raising the possibility of a common universal-like value across diverse neural systems. In future work, we will investigate whether such a universal-like value also emerges in other neural systems. More broadly, the competition-degree-based causal framework may provide a unifying generalized quantitative approach for understanding competitive harmony and its breakdown across diverse neural circuits and associated brain disorders.

## Appendix A: Izhikevich Neuron Models for CA1 Cells

In our CA1 SNN, we employ the Izhikevich spiking neuron model for CA1 cells [82–85] and adjust its intrinsic parameters for each cell type to match the corresponding experimentally obtained *f* - *I* curve [86–88].

The governing equations describing the time evolution of the dynamical states of individual cells in population *X* (*X* = PC, BC, or O-LM) are as follows:

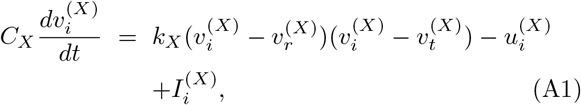

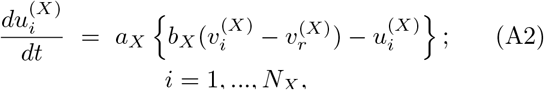

with the following after-spike reset rule:

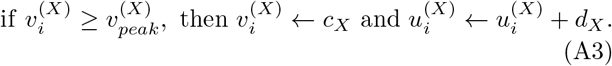

Here, *N*_*X*_ and 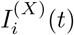 are the total number of cells and the current into the *i*th cell in population *X*, respectively. In Eqs. (A1) and (A2), the dynamical state of the *i*th cell in population *X* at time *t* (msec) is characterized by its membrane potential 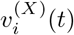 and the slow recovery variable 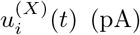. When 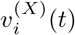 reaches the spike cutoff value 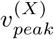, the cell fires a spike, after which 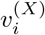 and 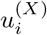 are reset according to Eq. (A3).

The Izhikevich neuron model for each population *X* has nine intrinsic parameters: *C*_*X*_ (pF): membrane capacitance, 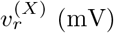: resting membrane potential,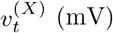: instantaneous threshold potential, *k*_*X*_ (nS/mV): parameter related to rheobase and input resistance, *a*_*X*_ (msec^*-*1^): reciprocal of the recovery time constant, *b*_*X*_ (nS): parameter controlling the voltage sensitivity of the recovery variable, *c*_*X*_ (mV): after-spike reset value of 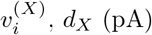 after-spike jump value of 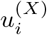, and 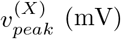 spike cutoff value. The values of these nine intrinsic parameters for the CA1 cells are listed in Table III.

The time evolution of 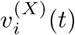 and 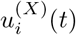 in Eqs. (A1) and (A2) is governed by the total input current 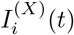 into the *i*th cell in population *X*:

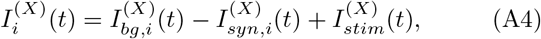

where 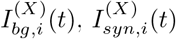, and 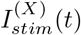 represent the background current from external regions not explicitly included in the model, the synaptic current, and the injected stimulation current, respectively. In our CA1 SNN, we set 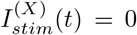 (i.e., no injected stimulation DC current).

The background current 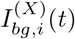 is modeled as the sum of 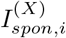 (spontaneous current responsible for spontaneous firing activity, corresponding to the time average of 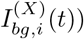 and 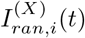 (random background current representing fluctuations around the time average of 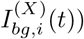.

For each CA1 cell population *X*, 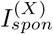 (independent of *i*) corresponds to the spontaneous in vivo current 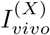 to reproduce the spontaneous in vivo firing rate 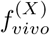 in the presence of synaptic inputs in the resting state (in vivo recordings in the awake resting state with tonic inputs from EC and CA3). The random background current 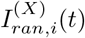 is given by:

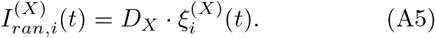

Here, *D*_*X*_ is the parameter controlling the noise intensity, and 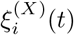 is Gaussian white noise with zero mean and unit variance [113–115]:

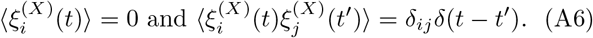

Table IV lists the spontaneous in-vivo currents 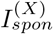, invivo firing rates 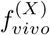, and random background noise intensities 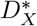 for CA1 cells in the awake resting state with tonic inputs from EC and CA3.

**TABLE IV:**
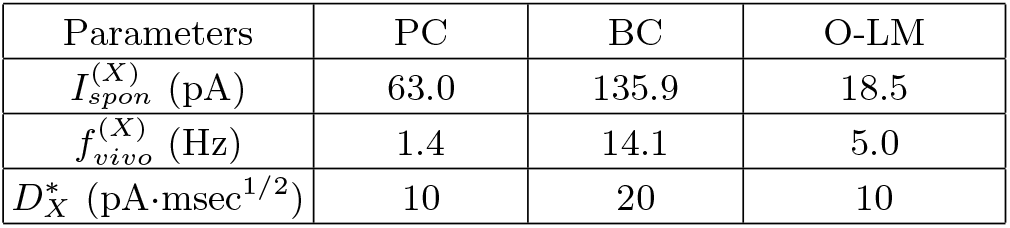
In vivo firing activities of hippocampal CA1 cells in the awake resting state with tonic external inputs from CA3 and EC (*f*_CA3_ = 1 Hz and *f*_EC_ = 3 Hz). Spontaneous current 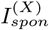, in vivo firing rate 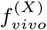, and random background noise intensity 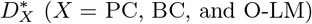.

| Parameters | PC | BC | O-LM |
|---|---|---|---|
| *I*<sup>(*X*)</sup><sub>*spon*</sub> (pA) | 63.0 | 135.9 | 18.5 |
| *f*<sup>(*X*)</sup><sub>*vivo*</sub> (Hz) | 1.4 | 14.1 | 5.0 |
| *D*<sup>\*</sup><sub>*X*</sub> (pA·msec<sup>1/2</sup>) | 10 | 20 | 10 |

## Appendix B: Excitatory and Inhibitory Synaptic Currents

We describe the excitatory and inhibitory synaptic currents 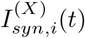 in Eq. (A4). There are two kinds of excitatory synaptic currents, 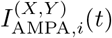 and 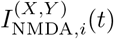, which are the AMPA (*α*-amino-3-hydroxy-5-methyl-4-isoxazolepropionic acid) receptor-mediated and NMDA (*N*-methyl-*D*-aspartate) receptor-mediated currents from the presynaptic source population *Y* to the *i*th cell in the postsynaptic target population *X*, respectively. The AMPA receptors mediate fast excitatory synaptic transmission, whereas NMDA receptors mediate slower excitatory synaptic currents with voltage-dependent Mg^2+^ block. In addition to these excitatory synaptic currents, we also consider two types of inhibitory synaptic currents, 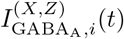 and 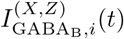, which are the GABA_A_ (*γ*-aminobutyric acid type A) receptor-mediated and GABA_B_ (*γ*-aminobutyric acid type B) receptor-mediated currents from the presynaptic source population *Z* to the *i*th cell in the postsynaptic target population *X*. The GABA_A_ receptors mediate fast, ionotropic inhibition, whereas GABA_B_ receptors generate slow, metabotropic inhibitory currents.

We follow the standard formalism for the synaptic currents, as in our previous works in the cerebellum [95, 96], the hippocampal dentate gyrus [97–100], and the basal ganglia [18, 19]. For receptor type *R* (*R* = AMPA, NMDA, GABA_A_, or GABA_B_), the synaptic current 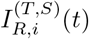 is given by:

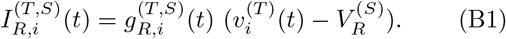

Here, 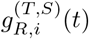 and 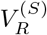 denote the synaptic conductance and synaptic reversal potential, respectively.

The synaptic conductance 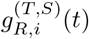 is given by:

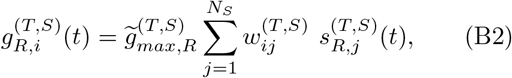

where 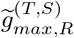 and *N*_*S*_ denote the maximum synaptic conductance and the number of cells in the source population *S*, respectively. The connection weight 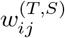 is 1 if the *j*th presynaptic cell is connected to the *i*th postsynaptic cell and 0 otherwise.

Here, 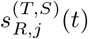 in Eq. (B2) represents the fraction of open postsynaptic ion channels opened by the binding of neurotransmitters released from the source population *S*. A sum of double-exponential functions 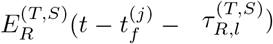 describes the time evolution of 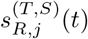 of the *j*th cell in the source population *S*:

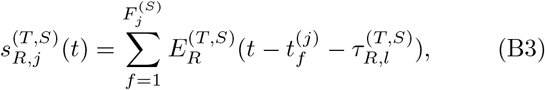

where 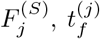, and 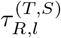 denote the total number of spikes of the *j*th cell, its *f* th spike time, and the synaptic latency time constant, respectively.

As in our previous studies in the hippocampal dentate gyrus [97–100], we use the following double-exponential function 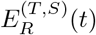:

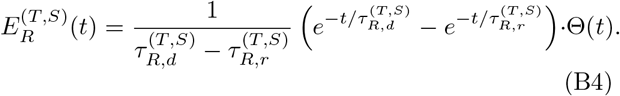

Here, Θ(*t*) is the Heaviside step function: Θ(*t*) = 1 for *t* ≥ 0 and 0 for *t <* 0. The parameters 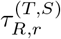 and 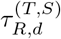 are the synaptic rise and decay time constants of the *R*-mediated synaptic current, respectively.

We also note that Mg^2+^ ions produce voltage-dependent blockade of postsynaptic NMDA channels. The fraction of unblocked NMDA channels is described by the sigmoidal function *f* (*v*^(*T*)^) [116, 117]:

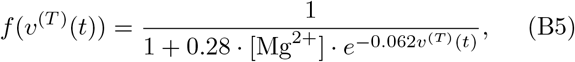

where *v*^(*T*)^ is the membrane potential of a cell in the target population *T* and [Mg^2+^] is the extracellular magnesium concentration ([Mg^2+^] = 1 mM). Then, the synaptic current into the *i*th cell in the target *X* population becomes

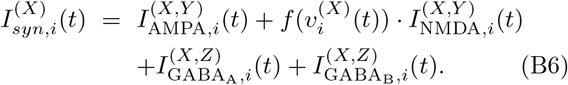

Table V shows the synaptic parameter values: *S*: source population, *T*: target population, *R*: receptor, 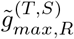: maximum synaptic conductance, 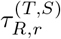: synaptic rise time constant, 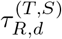: synaptic decay time constant, 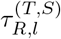: synaptic latency time constant, and 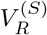: synaptic reversal potential.

**TABLE V:**
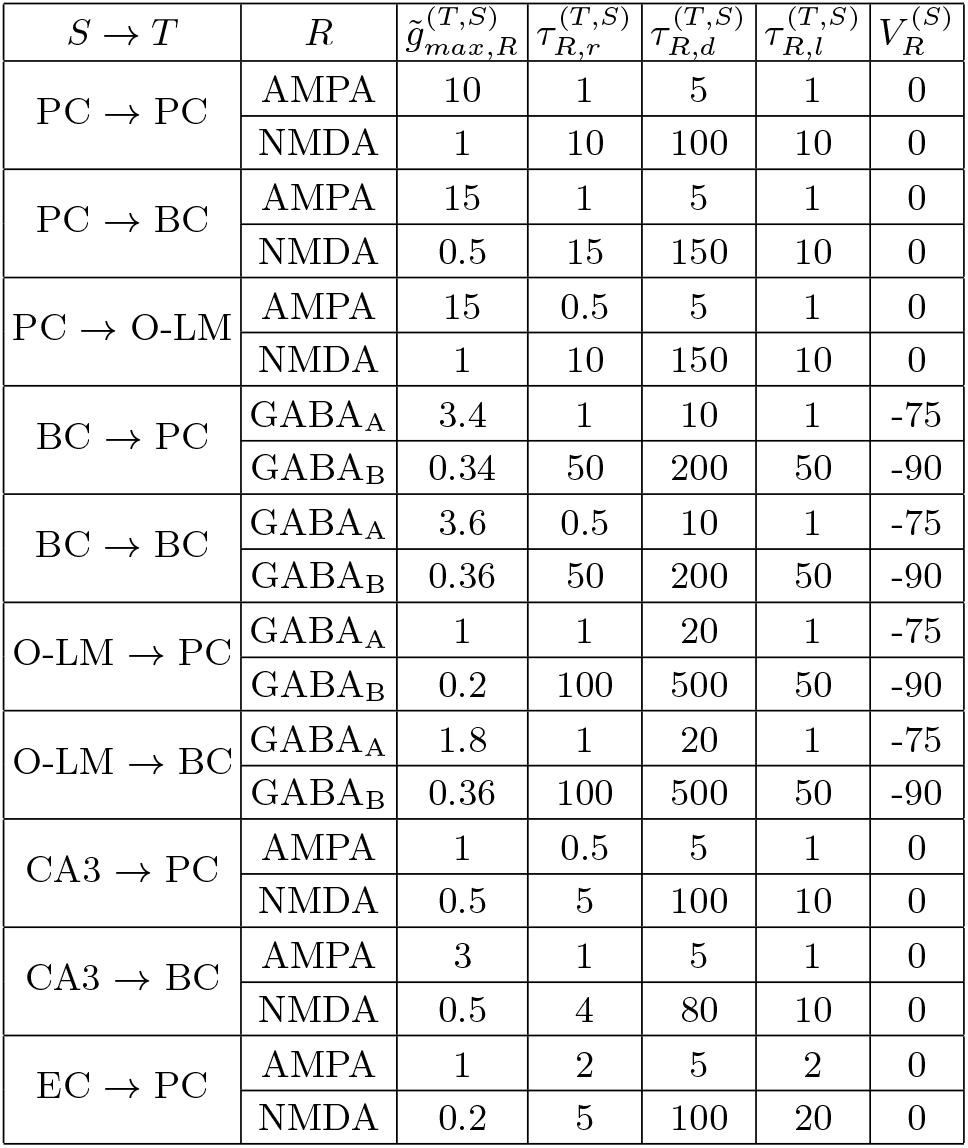
Synaptic parameter values. *S*: source population, *T*: target population, *R*: receptor, 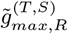: maximum synaptic conductance, 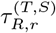: synaptic rise time constant, 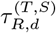: synaptic decay time constant, 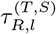: synaptic latency time constant, and 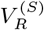: synaptic reversal potential.

| *S* → *T* | *R* | $\tilde{g}_{max,R}^{(T,S)}$ | *τ*<sub>*R,r*</sub><sup>(*T,S*)</sup> | *τ*<sub>*R,d*</sub><sup>(*T,S*)</sup> | *τ*<sub>*R,l*</sub><sup>(*T,S*)</sup> | *V*<sub>*R*</sub><sup>(*S*)</sup> |
|---|---|---|---|---|---|---|
| PC → PC | AMPA | 10 | 1 | 5 | 1 | 0 |
|  | NMDA | 1 | 10 | 100 | 10 | 0 |
| PC → BC | AMPA | 15 | 1 | 5 | 1 | 0 |
|  | NMDA | 0.5 | 15 | 150 | 10 | 0 |
| PC → O-LM | AMPA | 15 | 0.5 | 5 | 1 | 0 |
|  | NMDA | 1 | 10 | 150 | 10 | 0 |
| BC → PC | GABA<sub>A</sub> | 3.4 | 1 | 10 | 1 | -75 |
|  | GABA<sub>B</sub> | 0.34 | 50 | 200 | 50 | -90 |
| BC → BC | GABA<sub>A</sub> | 3.6 | 0.5 | 10 | 1 | -75 |
|  | GABA<sub>B</sub> | 0.36 | 50 | 200 | 50 | -90 |
| O-LM → PC | GABA<sub>A</sub> | 1 | 1 | 20 | 1 | -75 |
|  | GABA<sub>B</sub> | 0.2 | 100 | 500 | 50 | -90 |
| O-LM → BC | GABA<sub>A</sub> | 1.8 | 1 | 20 | 1 | -75 |
|  | GABA<sub>B</sub> | 0.36 | 100 | 500 | 50 | -90 |
| CA3 → PC | AMPA | 1 | 0.5 | 5 | 1 | 0 |
|  | NMDA | 0.5 | 5 | 100 | 10 | 0 |
| CA3 → BC | AMPA | 3 | 1 | 5 | 1 | 0 |
|  | NMDA | 0.5 | 4 | 80 | 10 | 0 |
| EC → PC | AMPA | 1 | 2 | 5 | 2 | 0 |
|  | NMDA | 0.2 | 5 | 100 | 20 | 0 |

## Appendix C: Output-Side Population and Individual Firing Activities of Interneurons

In this appendix, we study the output-side population and individual firing activities of inhibitory interneurons (BCs and O-LM cells) during epileptogenesis, complementing the analysis of PCs (principal excitatory neurons) in Sec. III. We consider two pathological mechanisms: axonal sprouting of PCs and loss of interneu-rons. Figure 10 illustrates the population firing activities of BCs and O-LM cells during the transitions from HP (green) to IIP (yellow) and then to IP (red).

**FIG. 10:**
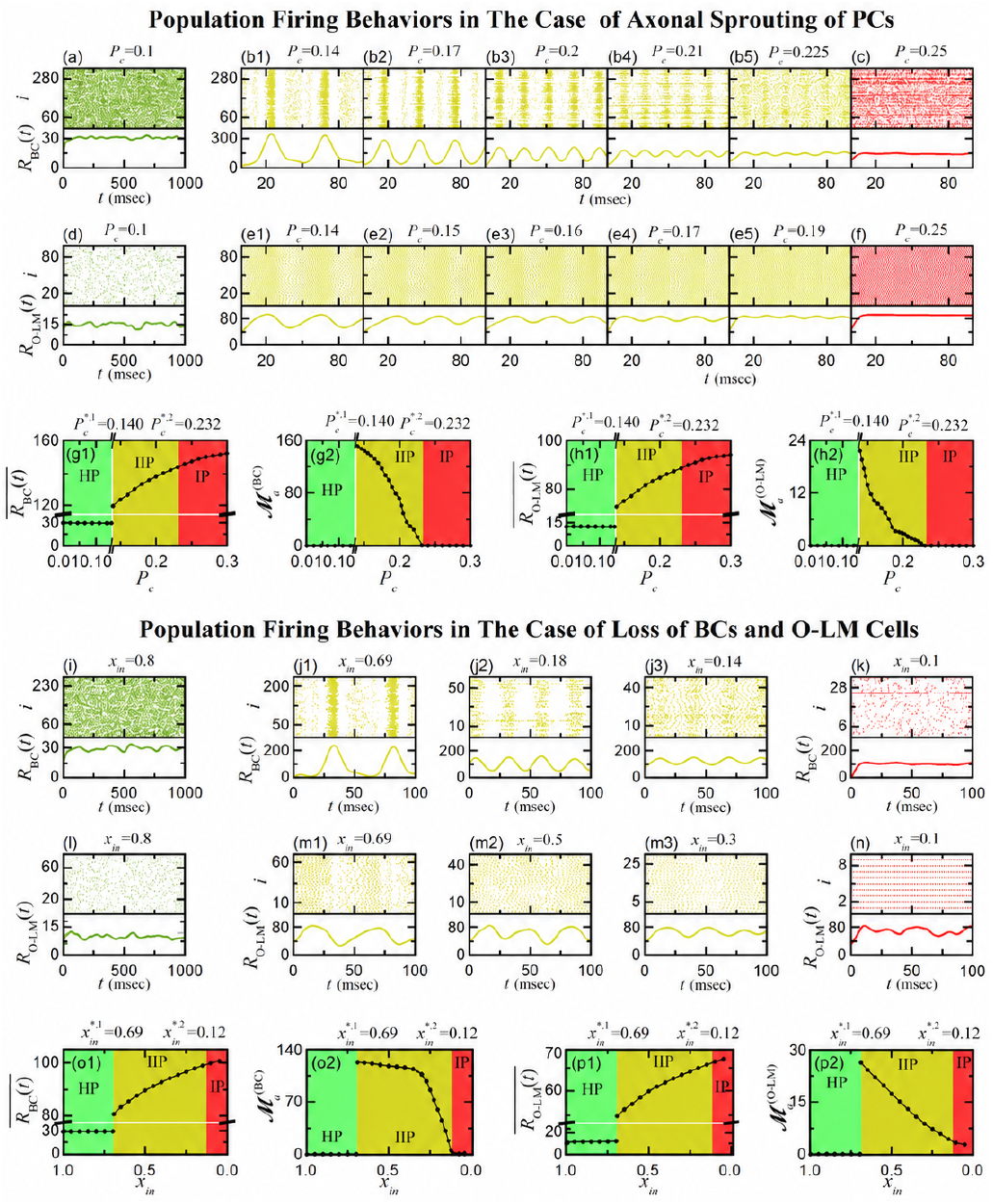
Output-side population firing activities of interneurons (BCs and O-LM cells) in the healthy and pathological states. Population firing activities in the case of axonal sprouting of PCs: (a)-(c) Raster plots of spikes and IPSRs *R*_BC_(*t*) of BCs. (d)-(f) Raster plots of spikes and IPSRs *R*_O*−*LM_(*t*) of O-LM cells. (g1) Plot of time-averaged IPSR 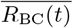 versus *P*_*c*._ (g2) Plot of amplitude measure 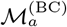 versus *P*_*c*._ (h1) Plot of time-averaged IPSR 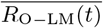 versus *P*_*c*._ (h2) Plot of amplitude measure 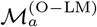 versus *P*_*c*._ Population firing activities in the case of loss of interneurons (BCs and O-LM cells) for *P*_*c*_ = 0.1: (i)-(k) Raster plots of spikes and IPSRs *R*_BC_(*t*) of BCs. (l)-(n) Raster plots of spikes and IPSRs *R*_O*−*LM_(*t*) of O-LM cells. (o1) Plot of time-averaged IPSR 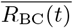 versus *x*_*in*_. (o2) Plot of amplitude measure 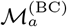 versus *x*_*in*_. (p1) Plot of time-averaged IPSR 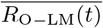 versus *x*_*in*_. (p2) Plot of amplitude measure 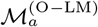 versus *x*_*in*_. Green, yellow, and red colors represent HP, IIP, and IP, respectively.

We first consider the effect of PC sprouting on the population activities of BCs and O-LM cells, which is well shown in Figs. 10(a)-10(h2). In the healthy phase (HP), interneuron spikes in their raster plots are randomly scattered, producing irregular IPSRs *R*_*X*_ (*t*) (*X* = BC and O-LM) without regular oscillations. The time-averaged IPSRs of BCs and O-LM cells are 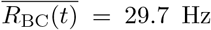 and 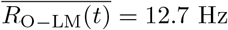 for *P*_*c*_ = 0.1, both higher than 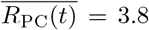 Hz of the PCs. The amplitude measures 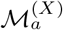, quantifying the degree of population synchronization, for *X* = BC and O-LM cells are nearly zero in HP, consistent with asynchronous firing.

At the 1st threshold of 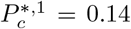, an abrupt transition to the interictal phase (IIP) occurs. At this point, narrower synchronized spike bands emerge for the BCs, while broader spike bands appear for the O-LM cells. Correspondingly, their IPSRs develop regular oscillations. At the transition from HP to IIP, both BCs and O-LM cells exhibit an abrupt increase in their time-averaged IPSRs (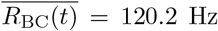 and 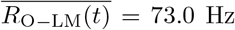) and in their amplitude measures (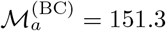 and 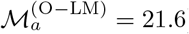).

Interestingly, at the threshold for the transition to IIP, their synchronization degrees differ: BCs show stronger synchronization than PCs 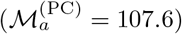, while O–LM cells remain less synchronized than PCs. In terms of time-averaged IPSRs, the hierarchy is 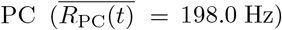 *>* BC *>* O–LM cell, in contrast to BC *>* O-LM *>* PC in HP.

These differences in the population synchronization degree *ℳ*_*a*_ can be understood in terms of synaptic connections among neuronal populations. BCs receive strong excitatory drive from PCs, but their activity is further shaped by dual inhibitory control: (i) mutual recurrent inhibition among BCs themselves and (ii) inhibitory input from O-LM cells. These two sources of inhibitory control promote phase-locking across the BC population and enhance their population synchronization. In contrast, O-LM cells receive excitatory inputs from PCs, but lack inhibitory regulation from BCs, which limits their ability to achieve strong population synchronization. This highlights the importance of synaptic connections, particularly inhibitory control, in shaping the different degrees of population synchronization of BCs and O-LM cells in IIP.

As *P*_*c*_ is further increased in IIP, spike bands approach each other, leading to a decrease in the synchronization degree (reflected by smaller oscillation amplitudes of the IPSRs). The synchronization degree of BCs (O-LM cells) 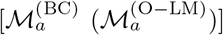 decreases more slowly (more rapidly) than that of PCs, as shown in the raster plots, IPSRs, and the corresponding amplitude measures *ℳ*_*a*_.

We also determine the population synchronization frequency 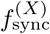 (*X* = BC and O-LM), corresponding to the oscillation frequency of the IPSR *R*_*X*_ (*t*). It is calculated as the inverse of the time-averaged oscillation period of *R*_*X*_ (*t*). As *P*_*c*_ increases in IIP, the population synchronization frequencies of both BCs and O-LM cells increase as adjacent spike bands approach one another in the raster plots of spikes. When comparing their IPSR oscillations, the population synchronization frequencies of BCs are comparable to those of PCs, whereas O-LM cells exhibit lower population synchronization frequencies. As in PCs, both BCs and O-LM cells are also found to exhibit dense population synchronization, characterized by multiple spiking of individual interneurons within each synchronized band. Consequently, the population synchronization frequency 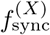 is lower than the population-averaged MFR 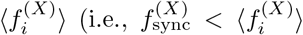; *X* = BC and O-LM). The population-averaged MFRs 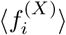 are presented below in the analysis of individual firing activities.

Finally, at the 2nd threshold 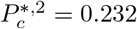, a transition to the ictal phase (IP) occurs via merging of spike bands. In IP, spikes of both interneuron types are randomly scattered, yielding irregular IPSRs *R*_*X*_ (*t*) (*X* = BC and O-LM) with high time-averaged values (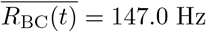 and 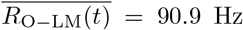 for *P*_*c*_ = 0.25). In IP, the amplitude measures 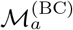 and 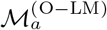 decrease to nearly zero, indicating that both interneuron populations become nearly desynchronized.

Next, we turn to the scenario of progressive interneuron loss, focusing again on the population firing activity of BCs and O–LM cells for *P*_*c*_ = 0.1. Figures 10(i)-10(p2) summarize how their population behaviors evolve as the surviving interneuron fraction *x*_*in*_ decreases, leading to transitions from HP to IIP and then to IP.

In the healthy phase (HP, *x*_*in*_ = 0.8), spikes of both BCs and O-LM cells are randomly scattered, yielding irregular IPSRs *R*_*X*_ (*t*) (*X* = BC and O-LM cells) without regular oscillations. The time-averaged IPSRs of BCs and O-LM cells are 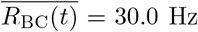 and 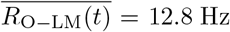 12.8 Hz, respectively, both again higher than that of PCs 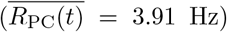. The amplitude measures 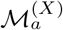 (*X* = BC and O-LM) vanish, indicating the absence of population synchronization in HP.

At the 1st threshold of 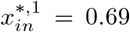, the system undergoes an abrupt transition into the interictal phase (IIP). At this point, narrower spike bands emerge in BCs, while broader spike bands appear in O-LM cells, and their IPSRs begin to oscillate regularly. At the transition from HP to IIP, their time-averaged IPSRs increase to 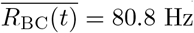 and 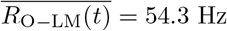, with amplitude measures 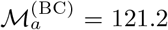 and 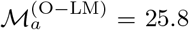, which quantify the degree of population synchronization. As in the sprouting case, BCs show stronger synchronization than PCs 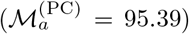, while O-LM cells remain less synchronized than PCs.

As interneuron loss progresses further within IIP (*x*_*in*_ *<* 0.69), the time-averaged IPSRs of both BCs and O–LM cells continue to increase due to hypoinhibition arising from interneuron loss. In contrast, the amplitude measures 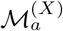 (*X* = BC and O-LM) progressively decrease, reflecting the weakening of collective synchronization as the inhibitory population shrinks. This decline results from the gradual approach of spike bands in the raster plots, which reduces the regularity and amplitude of IPSR oscillations, thereby weakening the degree of population synchronization.

We also determine the population synchronization frequency 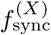 (*X* = BC and O-LM) in the same way as in the PC axonal sprouting case, namely, as the inverse of the time-averaged oscillation period of the IPSR *R*_*X*_ (*t*).

As interneuron loss progresses within IIP, the population synchronization frequencies of both BCs and O-LM cells increase because adjacent spike bands approach one another in the raster plots of spikes. Similar to the above case of PC axonal sprouting, in this case of interneuron loss, the population synchronization frequencies of BCs are also found to be comparable to those of PCs, whereas those of O-LM cells are found to be lower. As in the PC sprouting case, both BCs and O-LM cells also exhibit dense population synchronization with 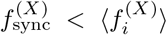 (*X* = BC and O-LM), due to multiple spiking of individual interneurons within each synchronized band.

At the 2nd threshold of 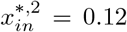, the system undergoes a transition to the ictal phase (IP). In this state, spikes of the BCs and O–LM cells become randomly scattered, and their IPSRs turn irregular. The time-averaged IPSRs are high (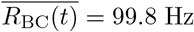 and 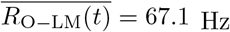 for *x*_*in*_ = 0.1), but the amplitude measures decrease to nearly zero, indicating that population synchronization is almost completely lost.

These observations motivate a comparison with PC sprouting, allowing us to examine whether similar interneuron dynamics and phase transitions occur under a different mechanism of PC hyperexcitability. Over-all, the population firing patterns of interneurons are similar in both scenarios. In both cases, hyperexcitable PCs drive interneuron activity, and reciprocal excitatory-inhibitory feedback loops between PCs and interneurons promote coordinated firing. The key difference lies in the mechanism of PC hyperexcitability: hyperexcitation via increased recurrent excitation in the sprouting case versus hypoinhibition in the interneuron loss case. Despite this difference, both cases exhibit a gradual approach of spike bands, decreasing amplitude measures, and an eventual transition to IP through the merging of spike bands. However, compared with PC axonal sprouting, the amplitude measures decrease more slowly during interneuron loss, indicating more persistent population synchronization of BCs and O–LM cells. We also note that BCs show higher synchronization than O–LM cells in both cases of PC sprouting and interneuron loss, owing to their stronger mutual interactions and inhibitory control.

Having analyzed the population-level firing behaviors of BCs and O-LM cells, we now examine their individual firing activities in both cases of PC sprouting and interneuron loss. We first consider the case of PC sprouting. Figures 11(a)–11(c) show the ISI histograms of BCs in the HP (green), IIP (yellow), and IP (red) states, while Figures 11(d)–11(f) display the corresponding histograms for O-LM cells. From these histograms, we compute the population-averaged ISI (*ISI*_*X*_) and the corresponding population-averaged MFR 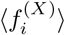, given by 1*/ ISI*_*X*_ (*X* = BC and O-LM), to characterize the average firing rates of individual interneurons. We then quantify the irregularity of individual firing using the coefficient of variation *CV*_*X*_.

**FIG. 11:**
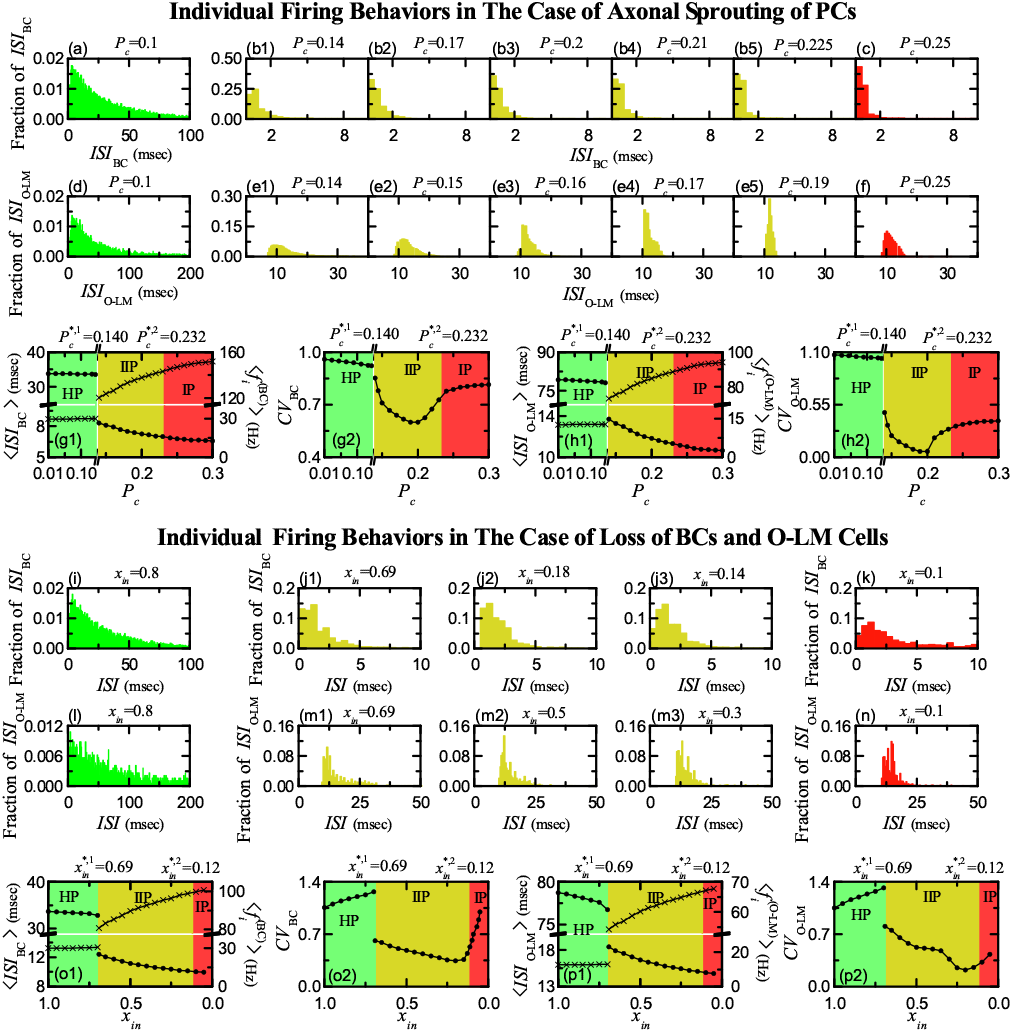
Output-side individual firing activities of interneurons (BCs and O-LM cells) in the healthy and pathological states. Individual firing activities in the case of axonal sprouting of PCs: (a)-(c) Population-averaged ISI histograms of BCs. (d)-(f) Population-averaged ISI histograms of O-LM cells. (g1) Plot of average ISI (*ISI*_BC_) of BCs (circles) and population-averaged MFR 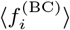 (= 1*/*(*ISI*_BC_)) (crosses) versus *P*_*c*_. (g2) Plot of *CV*_BC_ for BCs versus *P*_*c*_. (h1) Plot of average ISI (*ISI*_O*−*LM_) of O-LM cells (circles) and population-averaged MFR 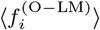 (= 1*/*(*ISI*_O*−*LM_)) (crosses) versus *P*_*c*_. (h2) Plot of *CV*_O*−*LM_ for O-LM cells versus *P*_*c*_. Individual firing activities in the case of loss of interneurons (BCs and O-LM cells) for *P*_*c*_ = 0.1: (i)-(k) Population-averaged ISI histograms of BCs. (l)-(n) Population-averaged ISI histograms of O-LM cells. (o1) Plot of average ISI (*ISI*_BC_) of BCs (circles) and population-averaged MFR 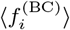 (= 1*/*(*ISI*_BC_)) (crosses) versus *x*_*in*_. (o2) Plot of *CV*_BC_ for BCs versus *x*_*in*_. (p1) Plot of average ISI (*ISI*_O*−*LM_) of O-LM cells (circles) and population-averaged MFR 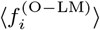 (= 1*/*(*ISI*_O*−*LM_)) (crosses) versus *x*_*in*_. (p2) Plot of *CV*_O*−*LM_ for O-LM cells versus *x*_*in*_. Green, yellow, and red colors represent HP, IIP, and IP, respectively.

During HP (green), the population-averaged ISIs 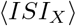 (circles) remain nearly constant at about 34 ms for BCs and 79 ms for O-LM cells. The corresponding population-averaged MFRs 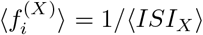 (crosses) also remain nearly constant at about 30 Hz for BCs and 13 Hz for O-LM cells, both higher than those of PCs. Their individual firing is irregular, as indicated by high coefficients of variation *CV*_*X*_ (*X* = BC and O-LM). At the 1st threshold of 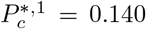, marking the transition to the interictal phase (IIP), both *ISI*_*X*_ and *CV*_*X*_ exhibit sudden drops, with the decreases being more pronounced for O-LM cells, while 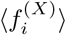 rises abruptly. In the subsequent IIP (yellow) and IP (red), (*ISI*_*X*_) (circles) decreases monotonically, while 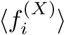 (crosses) increases monotonically. In contrast, *CV*_*X*_ displays a U-shaped profile: BCs reach a minimum *CV*_BC_ of 0.595 at *P*_*c*_ = 0.195, whereas O-LM cells reach a much lower minimum value of 0.049 at the same *P*_*c*_, indicating that O-LM cells fire much more regularly than BCs near their most ordered point. These variations highlight the distinct firing properties of BCs and O-LM cells. We also note that the population-averaged MFRs 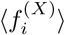, obtained from 1*/*(*ISI*_*X*_), are equivalent to the time-averaged IPSRs 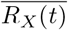 already shown in Fig. 10.

We next consider the case of interneuron loss at fixed *P*_*c*_ = 0.1. Figures 11(i)–11(k) show the ISI histograms of BCs, while Figs. 11(l)–11(n) display the corresponding histograms for O-LM cells across the three states of HP, IIP, and IP. As in the PC sprouting case, we analyze the population-averaged ISI (*ISI*_*X*_), the population-averaged MFR 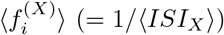, and the coefficient of variation *CV*_*X*_ (*X* = BC and O-LM).

In HP (green), the population-averaged ISIs ⟨*ISI*_*X*_ ⟩ remain nearly constant at about 33 ms for BCs and 78 ms for O-LM cells. The corresponding population-averaged MFRs 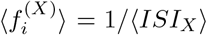 are also nearly constant at about 30 Hz for BCs and 13 Hz for O-LM cells (*X* = BC and O-LM). In contrast to the PC sprouting case, the coefficients of variation *CV*_*X*_ increase gradually and exceed 1, indicating progressively more irregular firing than a Poisson-like process, as in the case of PCs shown in Fig. 7(g). At the 1st threshold of 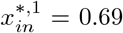, marking the transition to the interictal phase (IIP), both ⟨*ISI*_*X*_⟩ and *CV*_*X*_ exhibit sudden drops, while 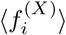 rises abruptly. During IIP (yellow) and IP (red), (*ISI*_*X*_) decreases monotonically, while 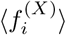 increases monotonically. In contrast, *CV*_*X*_ exhibits a U-shaped profile, similar to the PC sprouting case: BCs reach a minimum of 0.341 at *x*_*in*_ = 0.21, whereas O-LM cells reach a smaller minimum of 0.222 at *x*_*in*_ = 0.20, again indicating more regular firing of O-LM cells. These results show that, despite the different trends in *CV*_*X*_ during HP, the two pathological mechanisms produce qualitatively similar individual firing characteristics during IIP and IP.

## Acknowledgments

This work was supported by the National Research Foundation of Korea (NRF) grant funded by the Korea government (MSIT) (RS-2026-25586769).

## Author Contributions

S.-Y.K. conceived the study, developed the theoretical framework, and wrote the manuscript. W.L. performed simulations and generated figures. Both authors reviewed and approved the final manuscript.

